# Leaf ontogeny steers ethylene and auxin crosstalk to regulate leaf epinasty during waterlogging of tomato

**DOI:** 10.1101/2022.12.02.518836

**Authors:** B. Geldhof, J. Pattyn, P. Mohorović, K. Van den Broeck, V. Everaerts, O. Novák, B. Van de Poel

## Abstract

Developing leaves undergo a vast array of age-related changes as they mature. These include physiological, hormonal and morphological changes that determine their adaptation plasticity towards adverse conditions. Waterlogging induces leaf epinasty in tomato, and the magnitude of leaf bending is intricately related to the age-dependent cellular and hormonal response. We now show that ethylene, the master regulator of epinasty, is differentially regulated throughout leaf development, giving rise to age-dependent epinastic responses. Young leaves have a higher basal ethylene production, but are less responsive to waterlogging-induced epinasty, as they have a higher capacity to convert the root-borne and mobilized ACC into the inactive conjugate MACC. Ethylene stimulates cell elongation relatively more at the adaxial petiole side, by activating auxin biosynthesis and locally inhibiting its transport through PIN4 and PIN9 in older and mature leaves. As a result, auxins accumulate in the petiole base of these leaves and enforce partially irreversible epinastic bending upon waterlogging. Young leaves maintain their potential to transport auxins, both locally and through the vascular tissue, leading to enhanced flexibility to dampen the epinastic response and a faster upwards repositioning during reoxygenation. This mechanism also explains the observed reduction of epinasty during and its recovery after waterlogging in the *anthocyanin reduced* (*are*) and *Never ripe* (*Nr*) mutants, both characterized by higher auxin flow. Our work has demonstrated that waterlogging activates intricate hormonal crosstalk between ethylene and auxin, controlled in an age-dependent way.

## Introduction

Plants have developed several strategies to cope with low oxygen stress induced by flooding and submergence (Sasidharan et al., 2017). Morphological changes include escape responses such as hyponasty (Cox et al., 2003) and shoot elongation (Kende et al., 1998; Voesenek et al., 2004) to restore connectivity with the air. Other adaptations enhance the uptake and transport of oxygen through the formation of aerenchyma (Evans, 2004), adventitious roots (Vidoz et al., 2016) or hypertrophic lenticels (Shimamura et al., 2010). One of the more enigmatic responses caused by root anaerobiosis is the downward bending of leaves, called leaf epinasty. This nastic movement has been observed in several crops such as tomato (Jackson & Campbell, 1976), potato (Tonneijck et al., 1999), sunflower (Kawase, 1974) and cotton (Wiese & Devay, 1970). It is hypothesized that epinasty is an avoidance strategy to minimize intense solar radiation (Van Geest et al., 2012) and reduce plant transpiration (Else et al., 2009; Grichko & Glick, 2001).

A long time ago, exogenous ethylene was described to cause leaf epinasty in tomato (*Solanum lycopersicum*) (Doubt, 1917). Later, Jackson and Campbell (1976) established the relationship between waterlogging and an increase in ethylene production causing epinasty. Soon thereafter, it was shown that hypoxic conditions in the rooting zone induce both the production and transport of 1-aminocyclopropane-1-carboxylic acid (ACC), the immediate precursor of ethylene (Bradford & Yang, 1980). In tomato, ACC accumulates in hypoxic roots due to the upregulation of *ACC SYNTHASE7 (ACS7*) (Shiu et al., 1998), *ACS2* and *ACS3* (Olson et al., 1995). Subsequently, ACC is transported by the xylem (Bradford & Yang, 1980a) to the normoxic shoot, where it is converted into ethylene (English et al., 1995) by ACC OXIDASE1 (*ACO1*), causing the epinastic response (English et al., 1995).

Besides ethylene, other signaling molecules such as auxins (Keller & Van Volkenburgh, 1997; Sandalio et al., 2016), calcium (Lee et al., 2008) and radical oxygen species (ROS) (Pazmiño et al., 2014; Pazmiño et al., 2011) have been linked to epinastic movements of the leaf petiole and rachis. During leaf epinasty, both auxins (Lyon, 1963a, 1963b) and calcium are redistributed across the petiole (Lee et al., 2008), causing the cells on the adaxial side to elongate (Kazemi & Kefford, 1974; Ursin & Bradford, 1989a).

Early work has shown that auxins, besides ethyene, can evoke a strong epinastic response in tomato (Lyon, 1963). So far, it remains unknown if auxins act only through the induction of ethylene or not (Keller & Van Volkenburgh, 1997; Sandalio et al., 2016; Ursin & Bradford, 1989a). While auxins can induce epinasty in the presence of ethylene biosynthesis inhibitors (Keller & Van Volkenburgh, 1997) or in ethylene insensitive mutants (Romano et al., 1993), insensitivity to auxins in the auxin-resistant *diageotropica (dgt*) mutant does not impede epinasty after an ethylene treatment (Ursin & Bradford, 1989a). Although the interplay between both hormones is complex, ethylene has been described as a mediator of auxin biosynthesis, transport and signaling in other plant processes (Muday et al., 2012). SlIAA3, an Aux/IAA transcription factor, was identified as a possible point of crosstalk between auxin and ethylene signaling during leaf epinasty (Chaabouni et al., 2009). Moreover, both hormones have been associated with the production of ROS as a regulator of epinastic bending (Pazmiño et al., 2014).

Despite the identification of several signaling molecules involved in epinastic movements, their crosstalk and their action during flooding remain elusive. Furthermore, it is not known how leaves of different developmental stages respond to waterlogging and if there is an ontogenic hormonal regulation of leaf epinasty. In this study, we untangle the crosstalk between ethylene and auxin and assess how leaves of different ages respond to waterlogging and reoxygenation, especially in terms of epinasty.

## Materials and methods

### Plant material and growth conditions

Tomato (*Solanum lycopersicum*) seeds of the cultivar Ailsa Craig and the *Never ripe (Nr*) mutant were germinated in soil and later transferred to rockwool blocks. Seeds of the *SlPIN4-RNAi* and *pDR5::GUS* tomato lines were kindly provided by Prof. Carmen Catalá (Pattison & Catalá, 2012). The *are* mutant was kindly provided by Prof. Gloria Muday (Maloney et al., 2014). Tomato plants were grown in individual trays in a growth chamber or a greenhouse. The growth chamber had a temperature of 21 °C during the day and 18 °C during the night with a constant relative humidity of 65 %. Light was given in a 16 h/8 h day/night cycle using VS12 solar-spectrum mimicking LED lamps (Sunritec, China) with an intensity of 120 μmol s^-1^ m^-2^. In the greenhouse, the temperature was set at 18 °C during day and night with a humidity between 65 % and 70 %. In case the solar light intensity dropped below 250 W m^-2^ additional illumination was provided with high-pressure sodium lamps (SON-T). All plants received fertigation solution in their trays (growth chamber) or through drip irrigation (greenhouse).

### Waterlogging treatment

Tomato plants were grown until the eighth leaf stage. Subsequently, plants were transferred to individual trays filled with distilled water up to four cm above the rockwool surface to induce root hypoxia through natural oxygen consumption. During this period of waterlogging, the oxygen concentration in the root zone was monitored with an optical oxygen sensor (FireStingO_2_ Pyroscience). The hypoxia treatment started at 9 AM (1 – 2 h zeitgeber time) and was maintained for 24 to 96 h, after which the plants were removed from the trays to allow reoxygenation.

### Petiole morphology and epidermal anatomy

Leaf angles between the adaxial side of the petiole and the stem were measured with a protractor or a leaf angle sensor at the first two cm of the petiole base (described in Geldhof et al. (2021)). The length of petiole segments was measured between one and two centimeter marks at the petiole base. Subsequently, epidermal imprints were taken of the adaxial and abaxial side of the petioles after application of nail polish (Maybelline New York) for one hour. These imprints were visualized using an Olympus BX40 microscope and cell lengths were quantified in ImageJ.

### Plant and leaf physiology

Whole plant transpiration was measured using 24 digital lysimeters (KB 2400-2N scales, Kern) with a logging frequency of 10 s. Canopy cover changes were quantified through analysis of top view images (Nikon D3200) in ImageJ. Total plant CO_2_ consumption rate was monitored with SCD30 CO_2_ sensors (Sensirion) for individual plants in airtight PMMA boxes, flushed via an automated stop-and-flow system with a CO_2_ concentration between 400 and 600 ppm during the day. The maximum quantum yield of PSII (*F_v_/F_m_*) was measured on individual leaves with a continuous excitation chlorophyll fluorimeter (Handy Pea, Hansatech) after 30 min dark adaptation. Leaf photosynthesis, stomatal conductance and transpiration were determined for leaflet patches with an LCi compact portable photosynthesis system (ADC Bioscientific). Leaf fresh and dry weight were determined at different time points during waterlogging and subsequent reoxygenation.

### Ethylene measurements

Ethylene production of individual leaf petioles was determined by gas chromatography (GC-2010 Shimadzu). Two petiole sections of approximately 3 cm were sampled and incubated together in 5 mL cuvettes for 30 min (five replicates per treatment). Afterwards, the cuvettes were flushed to remove (wound) ethylene and sampled hourly following the same procedure. Ethylene measurements were performed for leaf 3, 5 and 7. Data was normalized for petiole fresh and dry weight.

### Histochemical staining of EBS::GUS and DR5::GUS

GUS activity was assessed following the protocol by Blakeslee and Murphy (2016). Petiole/stem segments of three cm were washed in buffer solution and vacuum infiltrated for 10 – 60 min in staining solution (1 mM X-Gluc; 100 mM sodium phosphate, pH 7; 10 mM EDTA; 0.5 mM potassium ferricyanide; 0.5 mM potassium ferrocyanide; 0.1 % Triton X-100), followed by 4 – 5 h (EBS::GUS) or overnight (DR5::GUS) incubation at 37 °C and clearing in 90 % ethanol:water (v/v). Petiole sections were visualized using an Olympus BX40 microscope and DR5::GUS staining was quantified by image analysis using a Python pipeline. Generation of the EBS::GUS reporter line is described in Supporting Information Method S1.

### 1-MCP and TIBA treatments

For the 1-methylcyclopropene (1-MCP) treatment, plants were transferred to airtight PMMA boxes the evening before the start of waterlogging. Each box contained three plants and half of the boxes were pre-treated with 1 ppm 1-MCP. For the 2,3,5-triiodobenzoic acid (TIBA) treatment, plants were sprayed with 1 mM TIBA and 0.05 % TWEEN-20 in water until runoff the evening before waterlogging. Control plants were sprayed with a 0.05 % TWEEN-20 water solution. Alternatively, TIBA was suspended in ethanol and lanolin (1 % m/m) and applied as a 0.5 – 1 cm ring around the leaf petiole.

### RNA extraction and RT-qPCR

Tomato leaves were snap frozen and ground in liquid nitrogen. RNA was extracted using the GeneJET Plant RNA Purification Mini Kit protocol (Thermo Scientific). DNA was removed using the RapidOut DNA Removal Kit (Thermo Scientific). Total RNA yield was assessed on the Nanodrop (Nanodrop Technology) and RNA quality was verified through gel electrophoresis. cDNA was synthesized using the iScript cDNA Synthesis Kit protocol (BIO-RAD). RT-qPCR was performed with a CFX96 (BIO-RAD) for 35 cycles. Four reference genes were selected based on earlier findings (Van de Poel et al., 2012) for normalization. Primers are listed in Supporting Information Table S1.

### ACC and MACC quantification

ACC and malonyl-ACC (MACC) content of different leaves were assessed based on the protocol described by Bulens et al. (2011) based on the Lizada & Yang (1979) method.

### ACO activity and abundance

ACO activity was derived from the *in vitro* ethylene production in airtight headspace vials, following Bulens et al. (2011). ACO was extracted using a modified protocol of Ververidis *et al.* (1991) as described by Mathooko *et al.* (1995) and Castellano *et al.* (2002). The ethylene content in the headspace was measured with a gas chromatograph (GC-2014 Shimadzu). Total ACO abundance was quantified using western-blotting according to Van de Poel et al. (2014), using a custom-made anti-ACO antibody targeting a conserved (ACO1-4) epitope.

### AMT activity

ACC-N-malonyl transferase (AMT) activity was derived from the *in vitro* MACC formation. AMT was extracted from snap-frozen crushed tissue (0.5 g) in 500 μL extraction buffer (200 mM Tris-Cl pH 8; 4 mM EDTA pH 8.5; 10 mM DTT; 2 mM benzamidine; 2 mM 6-amino-n-hexanoic acid; 0.5 mM N-α-p-tosyl-L-arginine methyl ester hydrochloride; 6 μM pepstatin; 2 μM leupeptin; 2 mM phenylmethanesulfonyl fluoride). After 30 minutes of incubation on ice and intermittent vortexing, the samples were centrifuged at 17.000 x g at 4 °C for 10 minutes. Next, 20 μL of the supernatant was added to 30 μL reaction mix (2 mM ACC; 0.5 mM malonyl-coA; 100 mM Tris-Cl pH 8; 1 mM DTT and 1 mM EDTA) and incubated for 3 h at 30 °C, followed by enzyme deactivation at 100 °C for 5 min. After removal of residual ACC by a cation exchanger (Dowex cation exchange column; 50WX8, 100-200 mesh), MACC was converted into ACC and subsequently converted into ethylene according to the protocol of Bulens et al. (2011).

### Auxin quantification

The auxin precursors tryptophan (TRP), tryptamine (TRA) and anthranilate (ANT), and IAA and its catabolite oxIAA were quantified in leaves and 2 cm petiole segments after 12 h, 24 h and 48 h of waterlogging (5 replicates per treatment). Leaves and petioles (leaf 1, 3, 5 and 7) were snap frozen, pulverized and analyzed following an UHPLC-ESI-MS/MS protocol described in Šimura et al. (2018).

### Statistical analysis

All analyses and visualizations were carried out in R. Treatments were compared using a Wilcoxon test for paired samples (α = 0.05) or a Dunn’s test with Bonferroni connection for multiple comparisons (α = 0.05). Empirical cumulative distributions (ECD) of cell lengths were compared using a Kolmogorov-Smirnov test (α = 0.05).

## Results

### Waterlogging-induced epinasty is leaf age-dependent

Tomato plants exposed to waterlogging show a quick epinastic bending of the leaves. By monitoring leaf angle dynamics in young tomato plants with 8 leaves, using a real-time clip-on IMU sensor (Geldhof et al., 2021), we observed an age-dependent (ontogenic) epinastic response towards waterlogging. The rapid decline of dissolved oxygen in the rooting zone (Figure 1A) caused more prominent downwards bending in older (leaf 3) and middle-aged (leaf 5) leaves, not recovering after 48 h of reoxygenation (Figure 1B & Supporting Information Fig. S1). Younger leaves (leaf 7) showed a similar epinastic response but were able to recover during reoxygenation.

**Figure 1:**
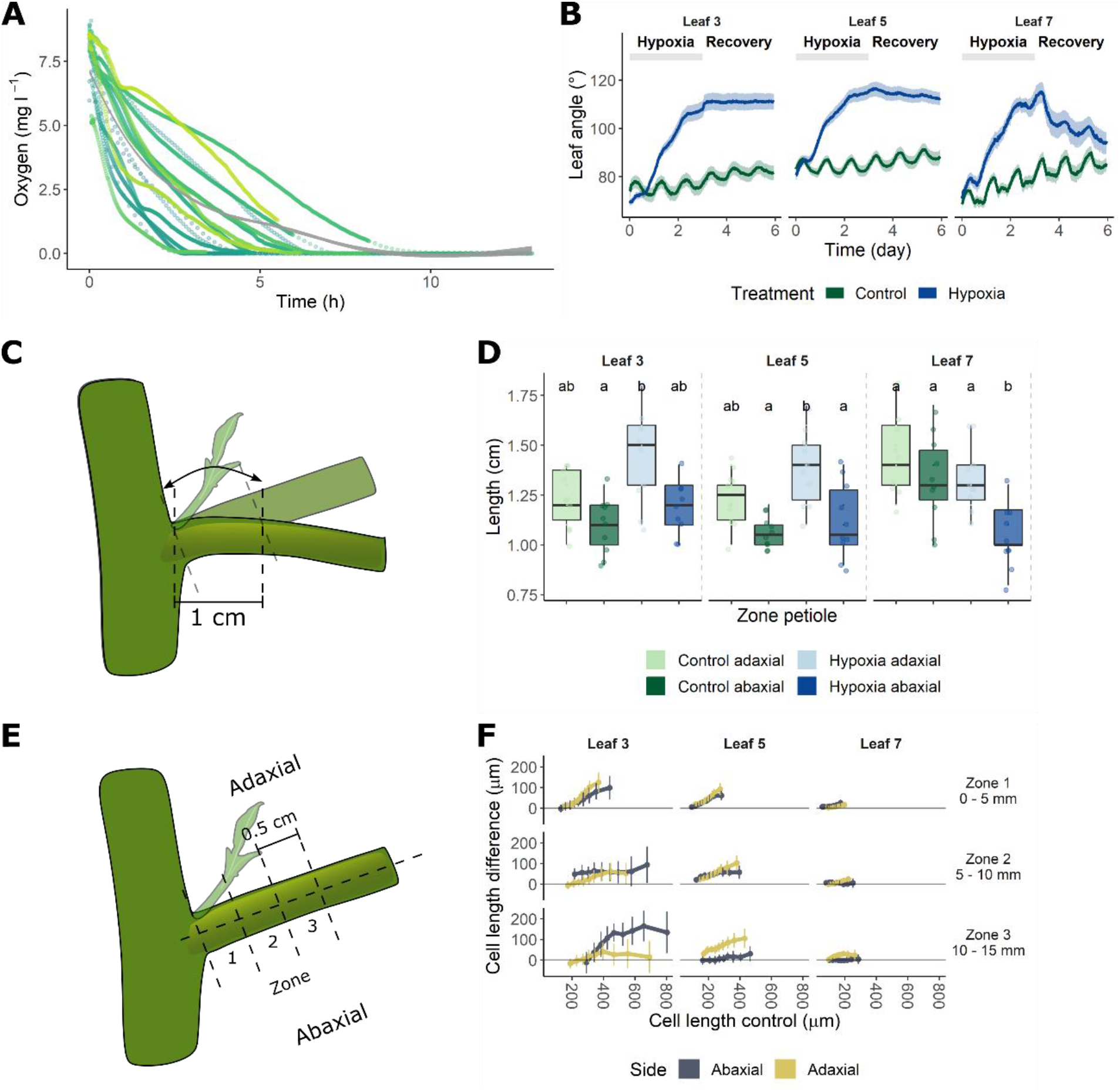
Ontogenic petiole angle dynamics, morphology and anatomy of tomato (Ailsa Craig; leaf 3, 5 and 7) after a waterlogging treatment. (A) Oxygen concentration in the rooting zone during the treatment, together with a smoothed model of the mean. Individual lines represent individual O_2_ consumption profiles. (B) Epinastic response of leaves of different ages during a 72 h waterlogging treatment and subsequent reoxygenation. (C – D) Length of the abaxial and adaxial sides (n = 10) of the first cm of the petiole after 48 h of treatment. (E – F) Differences in epidermal cell length distributions (n = 2 – 631 cells per treatment and zone for the adaxial and abaxial side per plant; 6 – 7 plants per treatment) between 48 h waterlogging and control conditions. The different zones in (E – F) refer to regions of the petiole of approximately 5 mm, starting at the petiole base (zone 1) and stretching out 15 mm along the petiole (zone 3). Significant differences are indicated with letters (α = 0.05).

To further characterize these epinastic movements at the cellular level, we compared the adaxial and abaxial petiole morphology and anatomy. Adaxial petiole segments of leaf 3 to 6 showed an enhanced elongation after 48 h of waterlogging, while this was seemingly reduced on the abaxial petiole side of young leaves (Figure 1C – D & Supporting Information Fig. S1). These variations in petiole elongation could explain the differences in angular dynamics between leaves. Subsequently, we examined cell dimensions using epidermal cell imprints. Both the adaxial and abaxial epidermal cell length increased after two days of root hypoxia, as shown by a significant shift in the empirical cumulative cell length distributions (ECD) (Figure 1E – G and Supporting Information Fig. S1). The ECD represents the proportion of cells smaller than or equal to a certain size (Supporting Information Method S3). We did not observe a difference in ECD for the adaxial side of leaf 8 and the abaxial sides of leaf 2, 7 and 8. In general, this shift suggests that elongation of cells is more pronounced at the abaxial side of the petiole, except for petiole segments located close to the stem (Figure 1E – F and Supporting Information Fig. S1).

To validate this enhanced abaxial elongation, we subsequently quantified the relative cell elongation during waterlogging by comparing cell length distribution quantiles in 0.5 cm zones on the adaxial and abaxial side of the petiole (Figure 1E). For a detailed description of this quantile shift method, we refer the reader to Supporting Information Method S3. In control conditions, cells at the abaxial side elongated relatively more than those at the adaxial side. During waterlogging, especially adaxial epidermal cells became proportionally larger, depending on the position along the petiole as well as leaf age. In addition, leaf development seemed to limit the extent of the differentiation between adaxial and abaxial epidermis cells. Altogether, epidermal cell elongation was respectively larger on the adaxial and smaller on the abaxial side of the petiole during waterlogging, causing downward bending of the petiole.

### Waterlogging reduces whole-plant transpiration and photosynthesis

To determine the effect of leaf epinasty on plant performance during waterlogging, we studied several physiological responses. Leaf epinasty significantly reduced canopy cover during waterlogging, only partly recovering after 7 days of reoxygenation (Figure 2A). Using real-time lysimetry, we observed that whole-plant transpiration rate dropped shortly after the start of the hypoxia treatment (31 – 47 % 1 day after treatment), and that it did not fully recover during reoxygenation (Figure 2B – C & Supporting Information Fig. S2). Accordingly, total plant CO_2_ uptake decreased during waterlogging and did not recover during reoxygenation (Figure 2D), indicating that both transpiration and photosynthesis are impaired at the onset of root hypoxia stress.

**Figure 2:**
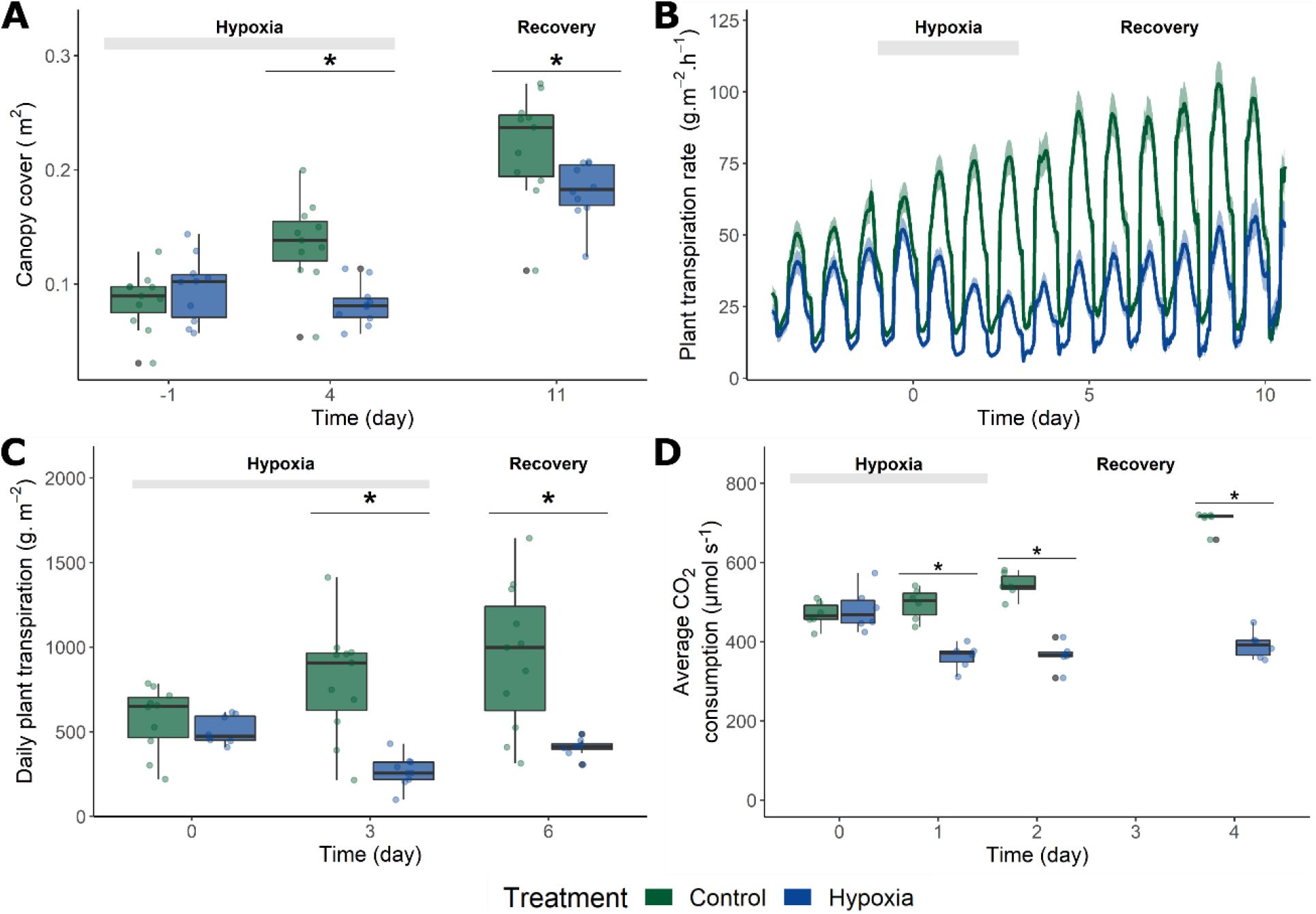
Physiological changes during waterlogging of tomato. (A) Canopy cover change determined from top view images (n = 10 – 11) after 96 h of waterlogging and 7 days of subsequent reoxygenation. (B) Total plant transpiration derived from real-time lysimetry (n = 10 – 11) before, during and after a waterlogging treatment and relative to the initial canopy cover. (C) Total daily plant transpiration integrated from (B). (D) Average daily plant CO_2_ uptake (n = 6) during and after a waterlogging treatment. Lines in (B) represent average transpiration rates +/− the confidence interval (90 %). Significant differences are indicated with an asterisk (α = 0.05).

### Leaf age determines physiological sensitivity towards root hypoxia

Given the effect of waterlogging on whole-plant performance, we wondered how leaves of different ages respond to waterlogging. Both leaf transpiration (Figure 3A) and leaf photosynthesis (Figure 3B) rate decreased throughout the treatment for all leaf age classes, but more rapidly in older leaves (Figure 3A – B). Recovery, especially for photosynthesis, was faster for young leaves during reoxygenation. The resilience of young leaves is not reflected by the plasticity of their stomatal conductance, which decreased drastically during waterlogging for all leaves and did not restore during reoxygenation. This suggests that recovery of photosynthesis in young leaves is largely independent of stomatal conductance (Figure 3C).

**Figure 3:**
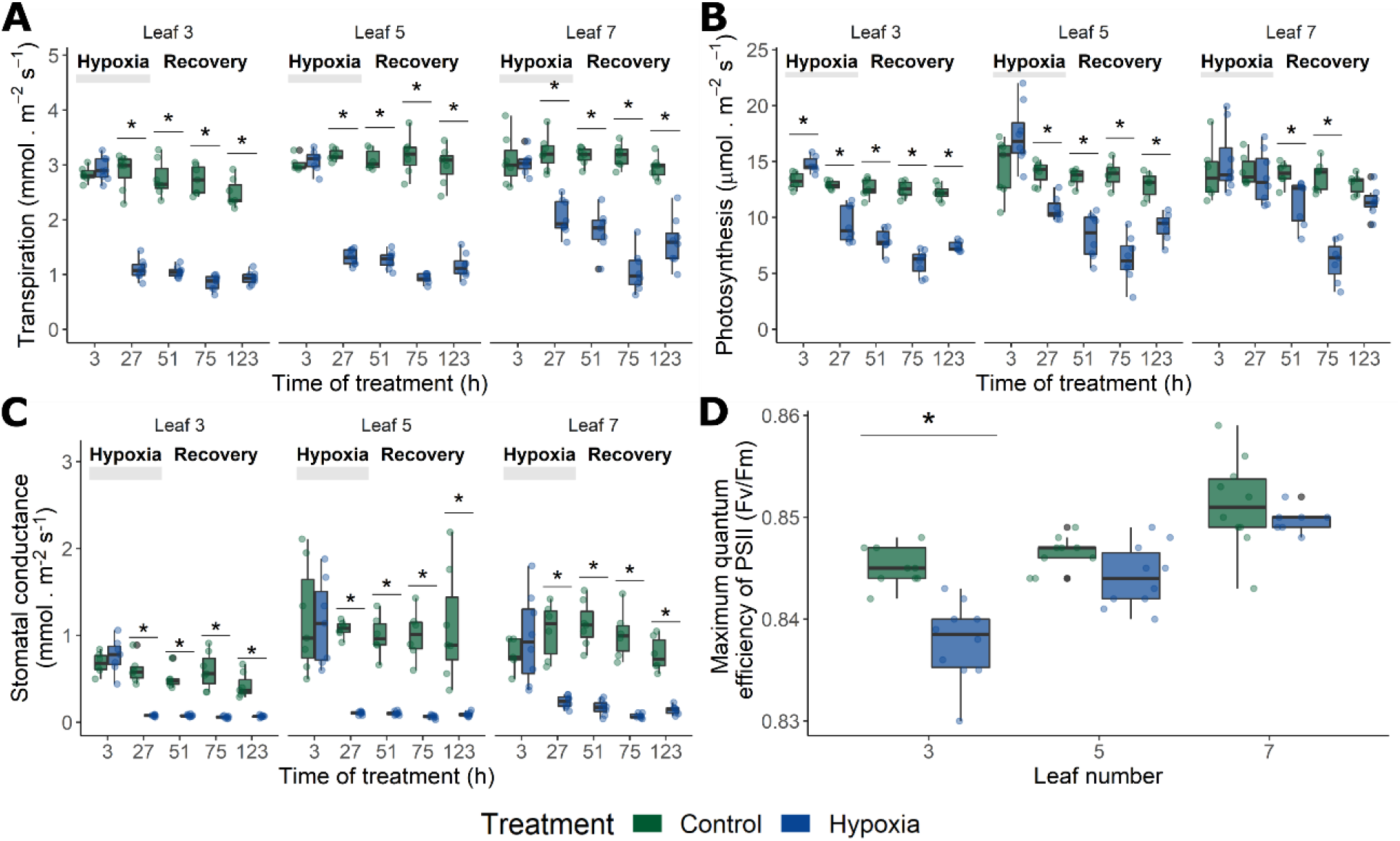
Leaf age-dependent changes of transpiration and photosynthesis during waterlogging and reoxygenation. Reduction of (A) leaf transpiration rate (B) leaf photosynthesis rate and (C) stomatal conductance, measured with an LCi photosynthesis system (n = 7 – 8). (D) Maximum quantum efficiency of PSII after 48 h of waterlogging treatment (n = 10). Significant differences are indicated with an asterisk (α = 0.05).

This discrepancy prompted us to quantify the maximum quantum efficiency of photosystem II (PSII; F_v_/F_m_), which decreased significantly in older (leaf 3) and middle-aged leaves (leaf 5) after 48 h and 54 h (data not shown) respectively, but not in young leaves (leaf 7; Figure 3D). Collectively, these leaf-specific results suggest that older leaves suffer more from waterlogging, while recently emerged leaves are able to limit root hypoxia induced PSII damage. Ultimately, reduced transpiration and carbon assimilation during 48 h of root hypoxia led to a decline in leaf fresh and dry weight mainly for middle-aged and young leaves, indicative of growth reduction (Supporting Information Fig. S3).

### Ethylene biosynthesis is ontogenically triggered by waterlogging

Ethylene plays a major role in the epinastic response during waterlogging, but the ontogenic regulation of the ethylene metabolism has not yet been explored. Therefore, we investigated ethylene biosynthesis of leaves of different ages during waterlogging. We first quantified ethylene production of leaf petioles of different ages (Figure 4A) and found that 24 h of waterlogging significantly enhanced ethylene production in petioles of all leaf ages, but the response was higher in older (leaf 3) and middle-aged (leaf 5) leaves.

**Figure 4:**
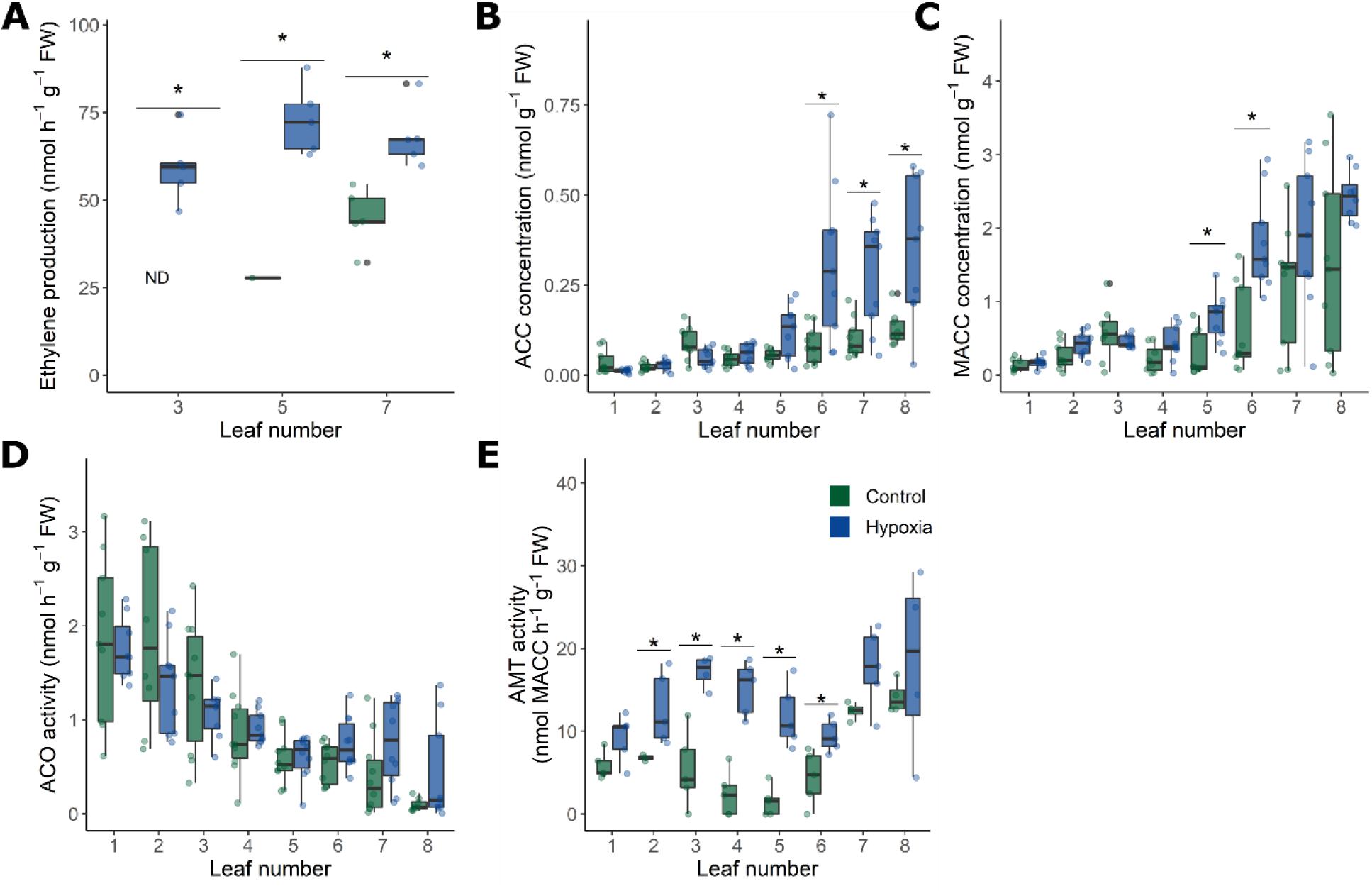
Ontogenic differentiation of ethylene biosynthesis during waterlogging. (A) Ethylene production of tomato petioles after 24 h of waterlogging (n = 5). (B – E) Changes in (B) the ethylene precursor ACC (n = 10), (C) the primary ACC conjugate MACC (n = 10), (D) enzyme activity of ACO (n = 10) and (E) enzyme activity of AMT (n = 5) after 24 h of waterlogging. Significant differences are indicated with an asterisk (α = 0.05).

To further unravel this ontogenic relation, several key intermediates of the ethylene biosynthesis pathway were measured for leaves of different ages during waterlogging. Our analysis showed that levels of the ethylene precursor ACC and its primary conjugate MACC were higher in younger leaves, and increased after 24 h of root hypoxia (Figure 4B & C). On the other hand, the total *in vitro* ACO activity was higher in older leaves, and did not change significantly after the hypoxia treatment (Figure 4D & E). The *in vitro* AMT activity, however, was stimulated in older and middle-aged leaves (leaf 3 – 5) by the hypoxia treatment, while it remained at a high level in young leaves (leaf 7). Together, these data indicate a complex ontogenic shift in the ACC metabolism. During waterlogging, ACC is transported from the roots to the shoot, and is most likely converted into ethylene in older leaves due to a high ACO activity. In contrast, the lower ACO and higher basal AMT activity in young leaves resulted in a lower ethylene production rate, leading to ACC accumulation and conversion into MACC. The increase of AMT activity of middle-aged leaves might limit ethylene production levels.

### Waterlogging induces differential regulation of the *ACO* gene family

In order to unravel the differences in ethylene production observed in leaves of different ages, we quantified total ACO protein abundance and the expression of several important *ACO* genes. Western blot analysis confirmed an ontogenic trend in ACO abundance, but did not show any increase after the hypoxia treatment (Figure 5A), matching the ACO *in vitro* activity (Figure 4D), with the exception of the oldest leaf (leaf 1), which had a high *in vitro* ACO activity and a low ACO protein abundance (Figure 4D & Figure 5A). Possibly, other ACO isoforms not detected by our antibody contribute to the ACO activity of leaf 1. A RT-qPCR analysis showed clear diversification in the expression of the *ACO* gene family with respect to ontogeny and hypoxia stress (Figure 5B – G). The expression of different *ACO*s under control conditions strongly depended on leaf age, with a slightly higher expression of *ACO2* and *ACO7* in younger untreated leaves. However, during root hypoxia, several *ACO* isogenes showed a strong upregulation, especially in older leaves for *ACO1, ACO2* and *ACO7* and in middle-aged leaves for *ACO5* (Figure 5B – G), not leading to an increase in ACO protein abundance (Figure 5A), but possibly contributing to the higher ethylene production rate (Figure 4A).

**Figure 5:**
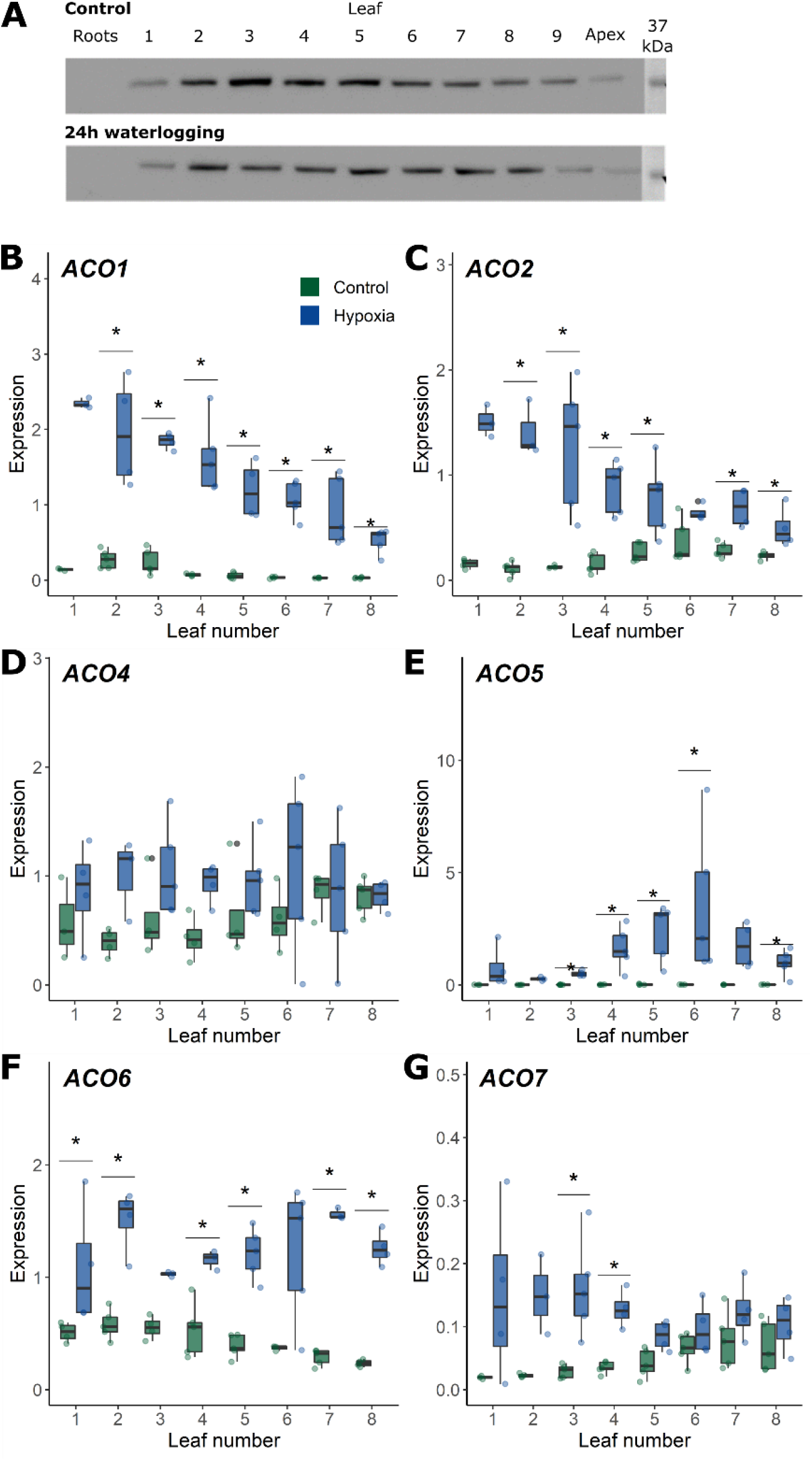
Effect of waterlogging on (A) the ACO abundance measured by western blotting (marker shows the 37 KDa protein size) and the relative expression of tomato (B) *ACO1,* (C) *ACO2,* (D) *ACO4,* (E) *ACO5,* (F) *ACO6* and (G) *ACO7* in leaves of different developmental stages and after 48 h of waterlogging (n = 5). Significant differences are indicated with an asterisk (α = 0.05).

### Ethylene sensitivity steers the epinastic response and its recovery

The ontogenic discrepancy between the ethylene production and its genetic and metabolic regulation at the leaf level prompted us to investigate the role of ontogeny in ethylene responsiveness. Although we did not observe waterlogging induced changes in an *EBS::GUS* reporter line, GUS levels seemed to change slightly during development. In general, the intensity of the GUS signal decreased in older petioles (Supporting Information Fig. S6), similar to petiole ethylene production levels (Figure 4A). Next, we evaluated ethylene sensitivity during waterlogging using a 1-MCP treatment and the *Nr* mutant, a natural mutant of ethylene receptor 3 (SlETR3) (Chen et al., 2019; Hackett et al., 2000; Lanahan et al., 1994). Our results showed that a 1-MCP pre-treatment strongly inhibited waterlogging-induced epinasty for leaves of all ages (Figure 6A). Dynamic leaf angle data of the *Nr* mutant supported these results, showing a strong reduction of the epinastic curvature during root hypoxia, especially for young leaves, followed by a faster recovery (Figure 6B & C). Altogether, these data confirm that ethylene signaling is key for epinasty during root hypoxia, but also reveal that ethylene sensitivity is differentially regulated in leaves of different ages.

**Figure 6:**
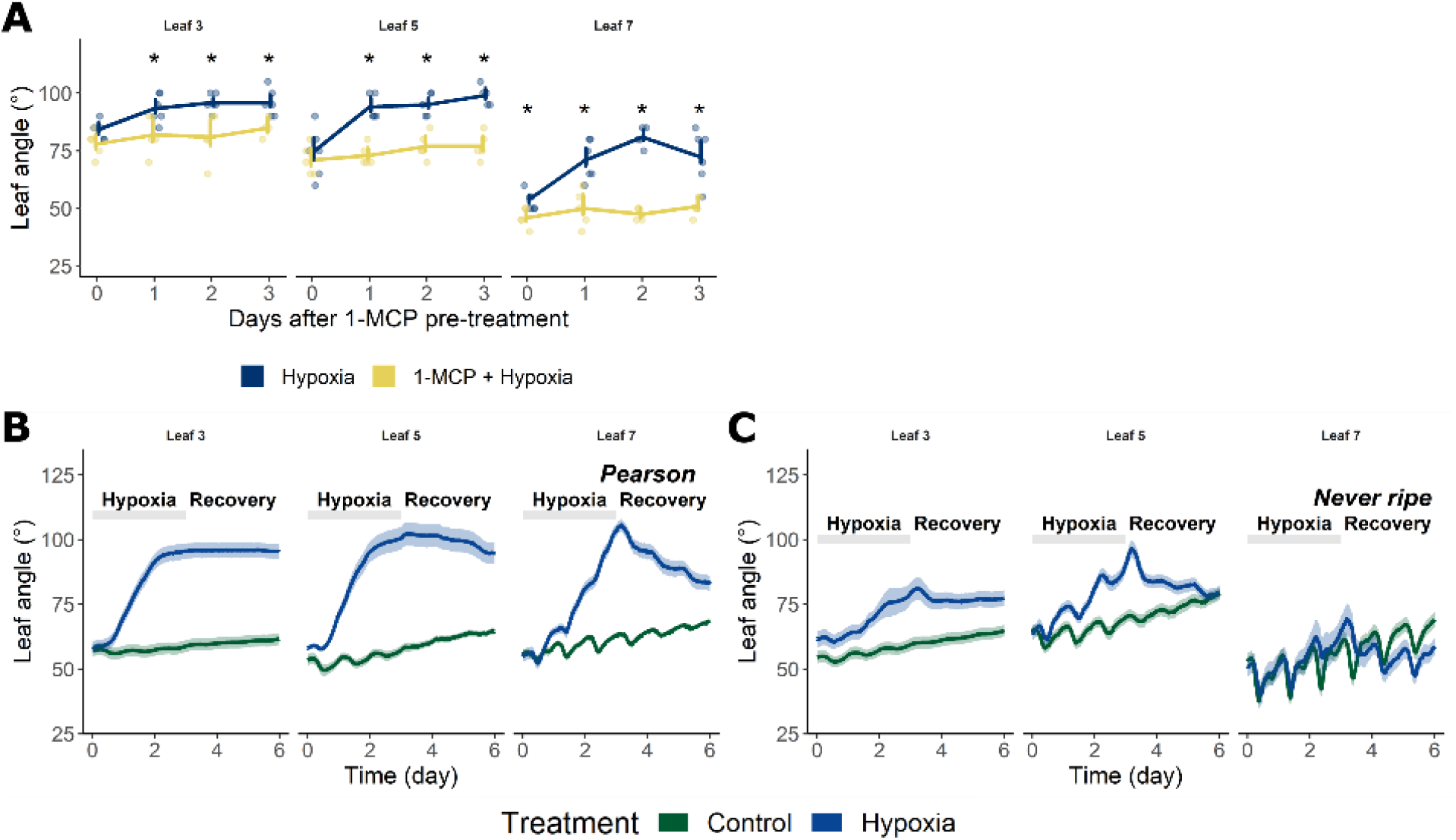
Ethylene sensitivity of leaves of different ages during waterlogging. (A) Effect of a 1-MCP (1 ppm) pre-treatment on the epinastic curvature after 48 h of waterlogging (n = 5 – 6). Significant differences are indicated with an asterisk (α = 0.05). (C – D) Epinastic response to waterlogging in the (B) Pearson background and (C) the *Nr* mutant. Lines in (C – D) represent average leaf angles +/– the confidence interval (90%).

### Waterlogging triggers auxin accumulation and responses in older leaves

Differentiation of ethylene biosynthesis and sensitivity might affect downstream regulation of epinasty through the auxin metabolism. To investigate auxin responses of petioles of different ages, we performed a histochemical analysis of a *DR5::GUS* reporter line (after 24 and 48 h of waterlogging) (Figure 7A & Supporting Information Fig. S7A). Auxin responses were generally more prevalent at the abaxial side of the petiole of old and mature leaves (leaf 3 – 5; Figure 7A), resembling the effect of an external ethylene treatment (Figure 7A). At the cellular level, GUS signals were stronger along the vascular tissue and the outer cell layers in petioles of young and middle-aged leaves under control conditions (Figure 7B – E). GUS staining in the parenchyma and collenchyma was also visible, but was mainly confined to abaxial cell layers of the petiole and the rachis during waterlogging (Figure 7D – E). These observations suggest that either local auxin production or its transport is augmented in the leaf petiole in an ontogenic way during waterlogging.

**Figure 7:**
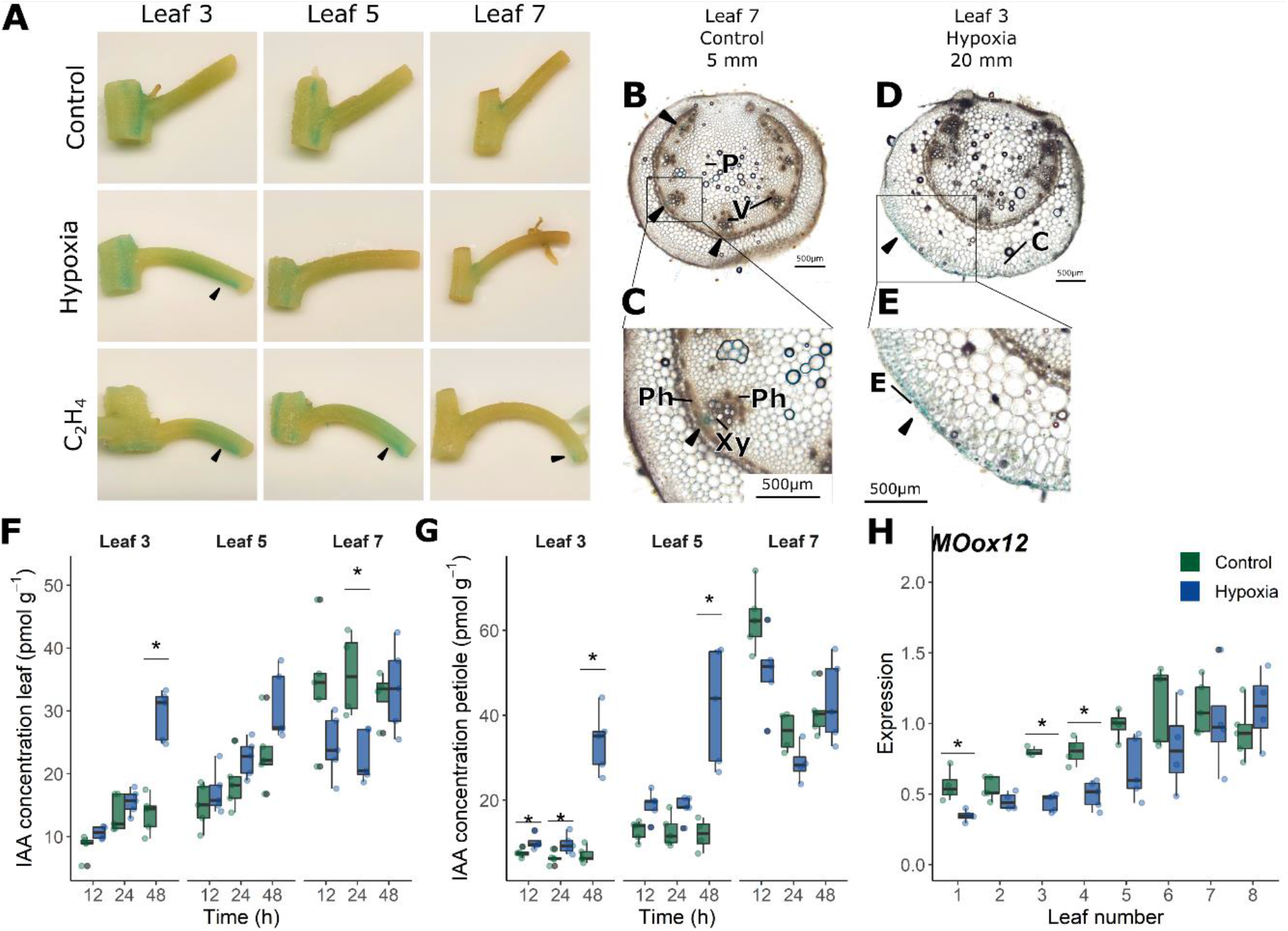
Auxin responses and metabolism in tomato leaves and petioles after a waterlogging treatment. (A) *DR5::GUS* expression in tomato petioles of leaf 3, 5 and 7 after 48 h of waterlogging or overnight ethylene treatment (10 ppm). (B – E) *DR5::GUS* expression in the petiole of (B – C) leaf 7 in control conditions, localized in the vascular tissue (5 mm from the proximal end) and (D – E) leaf 3 after 48 h of waterlogging, localized in the abaxial collenchyma (20 mm from the proximal end). Arrows indicate regions with GUS staining (V = vascular bundle; P = parenchyma; C = collenchyma; Ph = phloem; Xy = xylem; E = epidermis). (F – G) Auxin (IAA) levels in tomato (F) leaves and (G) petioles of different ages during the waterlogging treatment (n = 5). (H) Gene expression of an IAA biosynthesis gene *FLAVIN MONOOXYGENASE* in tomato leaves after 48 h of waterlogging (n = 5). Significant differences are indicated with an asterisk (α = 0.05).

Hormone analysis confirmed that IAA levels increased significantly in older and middle-aged leaves and petioles, especially after 48 h of waterlogging (Figure 7F – G & Supporting Information Fig. S7). Young leaves and petioles already contained a higher IAA level under control conditions, which remained stable (or even gradually decreased) during waterlogging. The auxin precursors tryptophan (TRP) and tryptamine (TRA) decreased rapidly (12 h) in all leaves during the waterlogging treatment, while anthranilate (ANT) levels only declined in young leaves (Supporting Information Fig. S7C – E). In petioles, the concentration of TRP and IAA increased after 48 h of hypoxia. Collectively, these data indicate that auxin production is dampened in young leaves but elevated in older petioles during waterlogging. The main auxin catabolite oxIAA increased significantly in petioles of all leaf ages, suggesting root hypoxia induces auxin conjugation (Supporting Information Fig. S7F). Both in control and waterlogged conditions, this conjugation was more prominent in young leaves. While IAA levels increased in older leaves and petioles, the expression of a *YUCCA-* like *FLAVIN MONOOXYGENASE* (Solyc08g068160), an IAA biosynthesis gene, was partially downregulated in old and middle-aged leaves after 48 h of waterlogging (Figure 7H). In young leaves, on the other hand, the expression of this auxin biosynthesis gene did not differ between the treatments, in accordance with the actual IAA levels. The time lag between the major shift in auxin metabolism (48 h after treatment) and the onset of the actual epinastic response (within 12 h) is indicative of a more complex regulation of auxin homeostasis, and could result from rapid changes in IAA mobility.

### Auxin transport is ontogenically gated during waterlogging

Auxin transport is known to regulate cellular development and elongation, but its role in the ontogenic control of leaf epinasty during waterlogging is unknown. A foliar treatment with TIBA, an inhibitor of polar auxin transport, resulted in a reduction of the epinastic response during waterlogging in older and middle-aged leaves (Supporting Information Fig. S8). However, the TIBA application itself also caused downwards bending and curling of control leaves. A local petiole application of TIBA lanolin paste confirmed that TIBA stimulates leaf epinasty, irrespective of the waterlogging treatment (Figure 8A). Furthermore, young leaves continued to bend down after the TIBA application, even during the reoxygenation phase. Intriguingly, epinastic bending of older and middle-aged leaves treated with TIBA stabilized during this phase. Besides the effect on leaf bending, inhibition of auxin export by TIBA also caused major leaf curling and changes in petiole morphology in young leaves, demonstrating the importance of polar auxin transport in leaf development (Supporting Information Fig. S9). Depending on leaf age, TIBA led to an accumulation of *DR5::GUS* expression upstream or downstream of the site of application of the lanolin paste. In young leaves, there was an increased GUS signal at the distal side, while in old leaves it was at the abaxial side of the petiole base (Supporting Information Fig. S10).

**Figure 8:**
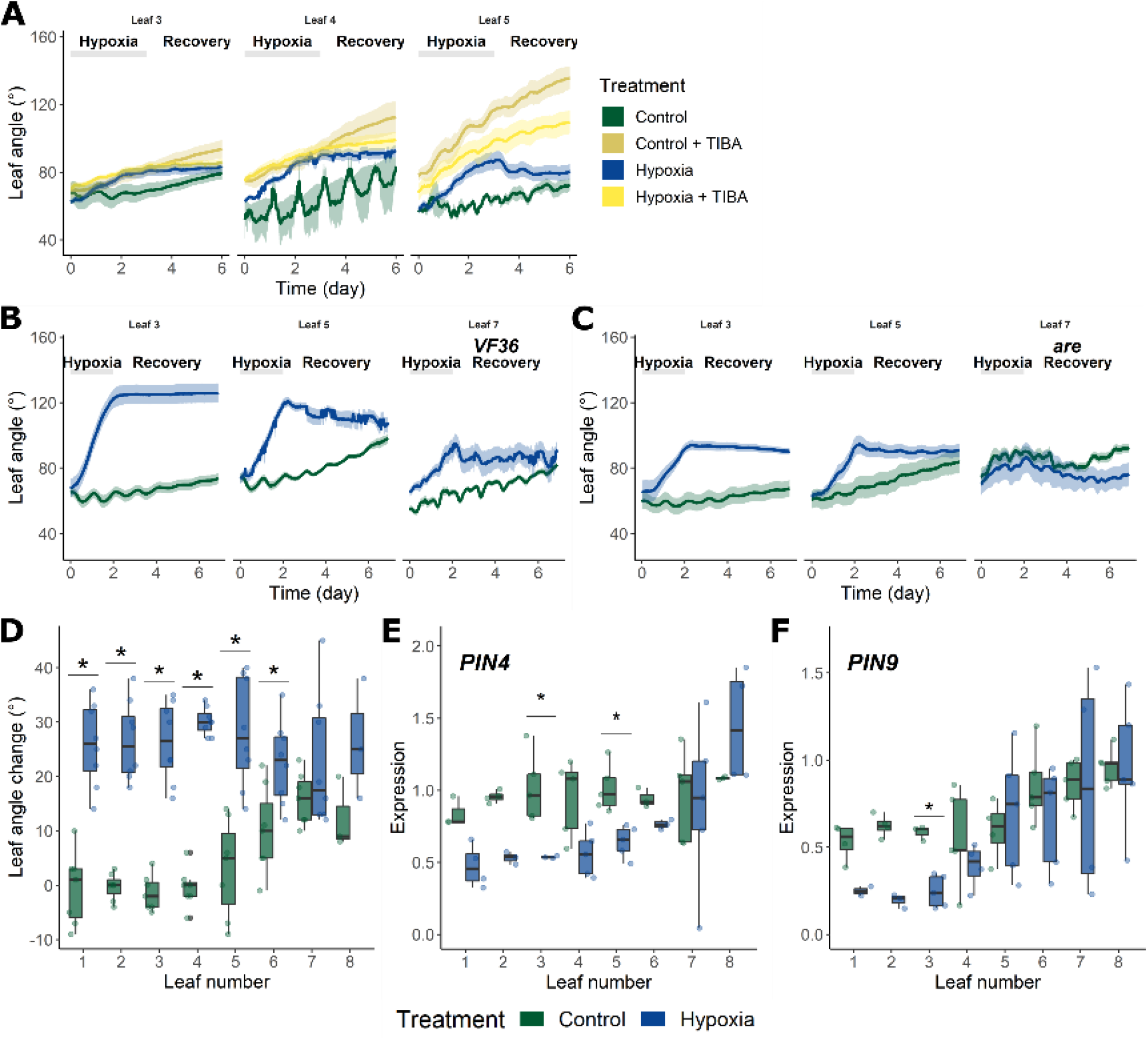
Ontogenic auxin transport in tomato leaves and petioles after a waterlogging treatment. (A) Effect of petiolar TIBA application (1 %) on leaf angle dynamics during waterlogging (n = 5 – 9). (B – C) Leaf angle change in the *are* mutant and its background VF36 during 48 h of waterlogging and reoxygenation (n = 5 – 10). (D) Epinastic response of a *PIN4-RNAi* line after 48 h of waterlogging (n = 7 – 8). (E – F) Gene expression of (E) *PIN4* and (F) *PIN9* transporters in tomato leaves after 48 h of waterlogging (n = 5). Lines in (D – E) represent average leaf angles +/− the 90 % confidence interval. Significant differences are indicated with an asterisk (α = 0.05).

The *are* mutant, characterized by a reduction of anthocyanins and their flavonol precursors, has a higher auxin flow (Maloney et al., 2014). This increased auxin flux in the *are* mutant reduced the intensity of the epinastic curvature (Figure 8B & C), corroborating the role of IAA transport in the waterlogging response. We evaluated the anthocyanin content in leaves, showing no alterations after 48 h of waterlogging (Supporting Information Method S2; Supporting Information Fig. S11). However, existing ontogenic differences might already interfere with auxin mobility.

Besides flavonols, auxin fluxes are predominantly controlled by PIN-formed (PIN) transporters. One of the exporters, PIN4, is strongly expressed in leaves (Pattison & Catalá, 2012). A *PIN4-RNAi* line showed a relatively normal epinastic movement during waterlogging (Figure 8D), but was already constitutively and mildly epinastic in old and mature leaves. These leaves also showed a decline in the expression of *PIN4* and another efflux carrier gene (*PIN9*) after 48 h of waterlogging (Figure 8E & F). Altogether, these results imply that auxin transport is impeded in old and mature leaves during waterlogging, in turn supporting IAA accumulation (Figure 7F) and signaling (Figure 7A) in these leaves.

## Discussion

### Leaf ontogeny defines epinastic curvature and recovery during waterlogging

Throughout development, leaves undergo several morphological, physiological and biochemical changes (Efroni et al., 2008). This process leads to the dynamic regulation of growth and determines the potential to respond to environmental cues (Rankenberg et al., 2021). We showed that one of these age-dependent responses is waterlogging-induced leaf epinasty, allowing young growing leaves to quickly re-adjust their posture after waterlogging.

This age-related plasticity has been described before for flooding-induced petiole elongation in *R. palustris* (Groeneveld & Voesenek, 2003), *R. pygmaeus* (Horton, 1992) and rice (*Oryza sativa*) (Alpuerto et al., 2022). In tomato, this age-related response seems to be regulated by cellular expansion during leaf growth (Figure 1). Waterlogging caused epinasty by reducing petiole and epidermal cell elongation of young leaves, mainly at the abaxial side, while promoting elongation in mature leaves, mainly at the adaxial side. This developmental dependency of cell and tissue elongation was also demonstrated for rice plants that need to reach a certain developmental stage before flooding can induce internode elongation (Ayano et al., 2014). In the monocot *Festuca arundinacea,* leaf elongation rates also increase with age (Xu et al., 2016).

### Young leaves retain a higher physiological plasticity during waterlogging

The effect of waterlogging on tomato physiology has been described without accounting for leaf ontogeny (Bradford & Hsiao, 1982; Else et al., 2009). We have shown that a short deprivation of oxygen in the root zone determines subsequent plant survival during reoxygenation (Figure 2), mainly depending on restoration of transpiration and photosynthesis in young leaves (Figure 3A – B). During drought stress in cotton (Jordan et al., 1975) and grapevine (Hopper et al., 2014), young leaves also showed higher resilience.

The plasticity of young leaves in terms of NPQ has been described as a possible protective mechanism against photoinhibition in Arabidopsis (Bielczynski et al., 2017). Young tomato leaves also seem capable of preventing these deleterious effects, as their PSII maximal quantum efficiency was unaffected by waterlogging in contrast to older leaves (Figure 3D). However, not every stress condition influences leaf performance in the same age-dependent way. Young expanding coffee leaves, for example, seem to be more prone to heat-induced reduction of both photosynthesis and stomatal closure (Marias et al., 2017).

### Leaf ontogeny defines the transcriptional regulation of *ACO* and the biosynthesis of ethylene

We have shown that both ethylene metabolism and *ACO* expression are ontogenically regulated in tomato (Figure 4 & Figure 5). Under control conditions, young tomato leaves harbor more ACC and produce a higher amount of ethylene, similar to young Arabidopsis leaves (Berens et al., 2019). Ontogenic regulation of *ACS* gene expression might contribute to this ACC gradient, as was shown before for maize (Young et al., 2004) and white clover (*Trifolium repens*) leaves (Murray & Mcmanus, 2005). During waterlogging, more ACC enters the leaves by xylem-mediated transport from the roots (Bradford & Yang, 1980a).

In white clover, ACC content and ethylene production of different leaves has been linked to differences in ACO activity (Chen & McManus, 2006; Hunter et al., 1999). In Arabidopsis, *AtACO1* is upregulated in roots during short term waterlogging (Hsu et al., 2011), and in both roots and shoot during long term waterlogging (Ventura et al., 2020) or hypoxia (Hsu et al., 2011). Our own transcriptional analysis showed that *SlACO5,* belonging to the same clade as *AtACO1* (Houben & Van de Poel, 2019), is strongly upregulated in an ontogenic way, with the highest expression in mature and younger leaves (Figure 5). The closest ortholog to *AtACO5,* the only ACO previously linked to leaf bending (Rauf et al., 2013), is *SlACO7* (Houben & Van de Poel, 2019) and is predominantly upregulated in older and mature leaves. This also counts for *SlACO1,* explaining the reduced epinastic bending in older leaves of its antisense line during waterlogging in tomato (English et al., 1995). The differential expression of *ACO* genes by age and waterlogging does not reflect in differences in ACO protein abundance nor *in vitro* activity, hinting towards a possible post-translational regulation ACO.

This indicates that differentiation of the ACOs provides a mechanism to control ethylene-regulated stress responses in a tissue- and age-specific way (Figure 9). In tomato, this leads to higher ethylene production in younger leaves during control and waterlogging conditions (Jackson & Campbell, 1976), but not during drought stress (Abou Hadid et al., 1986). In petioles, the increased ethylene production is relatively higher for older and mature leaves (Figure 4A). One mechanism that could further contribute is conjugation of ACC (Pattyn et al., 2021). Indeed, MACC accumulation is higher in young leaves, probably through higher basal AMT activity (Figure 4E). In contrast, old leaves activate AMT during waterlogging without a detectable change in MACC content.

**Figure 9:**
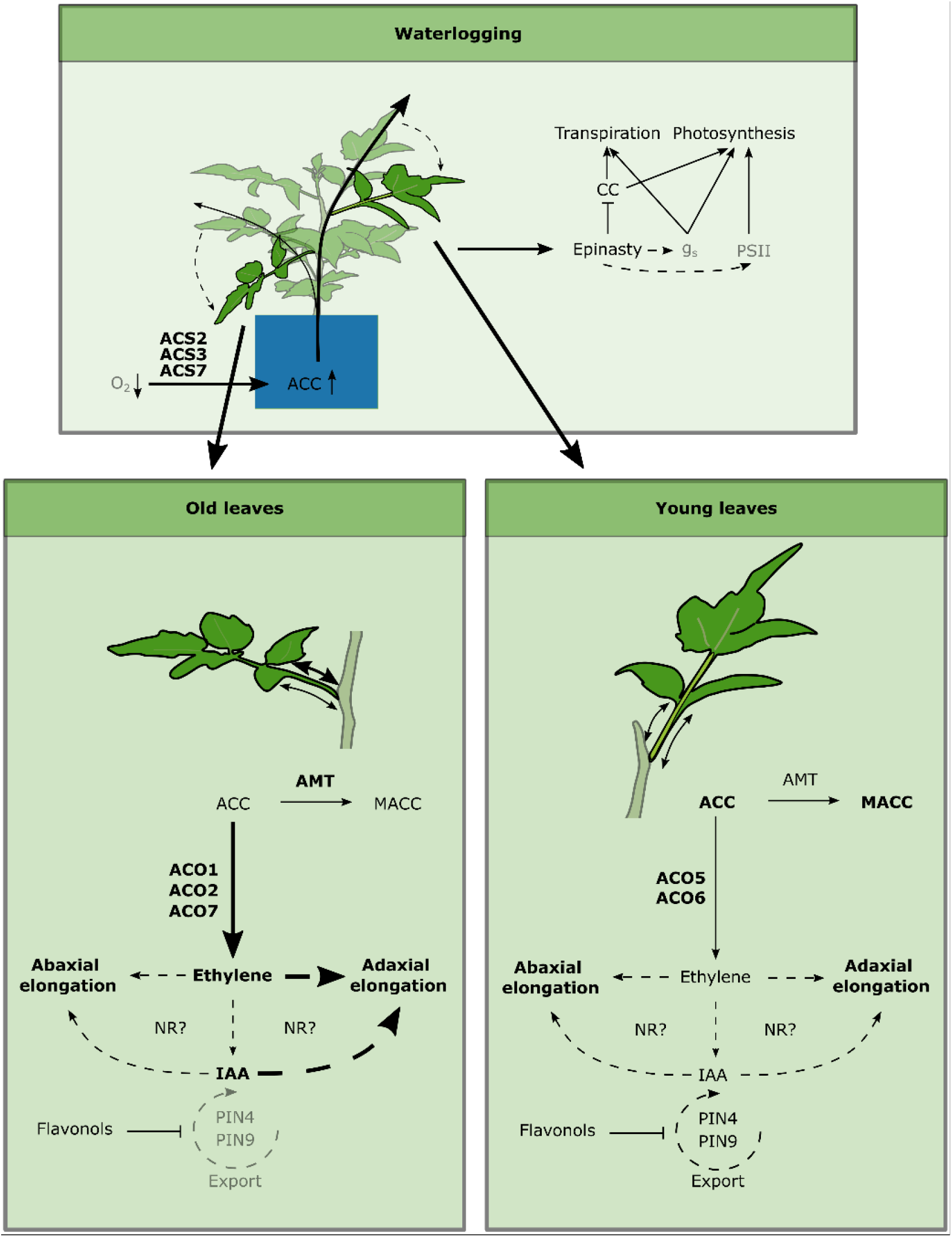
Schematic overview of the ontogenic differentiation of the epinastic response during waterlogging in tomato. Low oxygen conditions in the roots stimulate ACC production, at least by ACS2, ACS3 and ACS7 (Olson et al., 1995; Shiu et al., 1998), leading to accumulation of ACC in the root zone. This ACC is transported to the shoot, where it is distributed mainly towards young sink leaves, capable of effectively conjugating ACC into the inactive MACC. Differential expression of the *ACO* gene family allows for intricate regulation of local ethylene production, which is stimulated mainly in older leaves. Ethylene itself activates cellular elongation, in part through NR signaling and the regulation of local auxin levels. Auxin export is inhibited in old leaves by reduced expression of *PIN4* and *PIN9* auxin exporters, promoting its accumulation at the base of the petiole. This in turn disrupts local elongation dynamics within the petiole, leading to a relatively higher adaxial cell elongation and ultimately epinastic bending (and reduction in canopy cover, CC). In young leaves, auxin export capacity is sustained and hormonal balances are faster restored, leading to enhanced morphological plasticity and resilience during waterlogging and recovery. Most likely, other factors come into play to define the proper transduction of polar responsiveness within the petiole, as inhibition of auxin export from the leaf alone (by TIBA) evokes specific secondary and age-related responses regarding leaf positioning, independent of waterlogging-induced epinasty.

### Leaf plasticity is determined by ontogenic changes in ethylene sensitivity

Leaf plasticity towards stress can also be explained by developmental differences in ethylene sensitivity and signaling (Edelman & Jones, 2014; Kanojia et al., 2020; Kim et al., 2009; Li et al., 2013). In submergence-tolerant rice, ethylene sensitivity is developmentally controlled by differential regulation of *SUBMERGENCE1A (SUB1A*) dependent pathways, leading to a stronger growth reduction in young leaves (Alpuerto et al., 2022). In contrast, young rosette leaves of Arabidopsis seem to be more hyponastic after application of exogenous ethylene (Vandenbussche et al., 2003) and during waterlogging (Rauf et al., 2013). Similar differences in ethylene sensitivity could dampen leaf epinasty in young leaves during waterlogging (Figure 1B). This age-dependent sensitivity to ethylene and the corresponding epinastic response have also been reported for other *Solanaceae* species (Edelman & Jones, 2014).

A 1-MCP treatment prevented waterlogging-induced epinastic bending of all leaves (Figure 6A), indicating that ethylene signaling is crucial for epinastic bending (Figure 9). The *Nr* mutation also inhibits epinastic responses (Barry et al., 2005; Lanahan et al., 1994). The residual epinastic curvature observed in older and mature leaves of the *Nr* mutant during waterlogging (Figure 6C – D) indicates that the ethylene signal still gets partially transduced by other receptors (Chen et al., 2019; Hackett et al., 2000; Lanahan et al., 1994).

### Elevated auxin levels in mature tomato leaves contribute to epinasty during waterlogging

Ethylene sensitivity is often intertwined with auxin homeostasis, signaling and transport (Muday et al., 2012), which is developmentally regulated in Arabidopsis (Ljung et al., 2002). In general, young leaves are biosynthetically active and rely on different modes of auxin transport, leading to ontogenic differences in auxin sensitivity (Ljung et al., 2002). As a result, both auxin depletion and accumulation might yield leaf-specific responses. The reduction of auxin levels during tomato leaf ageing (Figure 7F – G) could explain developmental differences in leaf bending (Figure 1 D&F). However, the role of auxins in developmentally steered elongation might be more complex. In rice, for example, submergence-induced leaf elongation is less prominent in mature leaves, despite the higher accumulation of auxins (Alpuerto et al., 2022). On the other hand, rice coleoptile elongation under submergence is directly related to an increased auxin content (Nghi et al., 2021).

In tomato, waterlogging elevates auxin levels in mature leaves and decreases auxin precursors (TRP and TRA) in older leaves (Figure 7F – G; Supporting Information Fig. S7). This indicates there is enhanced IAA synthesis during waterlogging, despite one *YUCCA* homolog not being upregulated (Figure 7H). Another mechanism that might contribute to differential age-related auxin levels is conjugation. In Arabidopsis, the glucosyltransferase *UGT74D1* is mainly expressed in young leaves, and overexpression of *UGT74D1* results in increased IAA levels and ultimately in less erect leaves (Jin et al., 2021). Oxidation of IAA into the inactive oxIAA in young tomato petioles (Supporting Information Fig. S7) might provide an additional feedback mechanism to control auxin homeostasis.

Besides IAA synthesis and breakdown, reduced auxin export by PIN transporters, reflected by a reduced expression of *PIN4* and *PIN9* (Figure 8E – F), possibly contributes to IAA accumulation in older leaves during waterlogging (Figure 7F – G; Figure 9). Accumulation of auxins due to tissue-specific inhibition of IAA transport seems likely, as the *PIN4* RNAi line (Figure 8D) and an application of the auxin export inhibitor TIBA (Figure 8A) both showed epinasty, irrespective of a waterlogging treatment and leaf age. A similar mechanism has been described in the *Atmdr1* mutant of Arabidopsis, characterized by inhibition of auxin transport and leaf epinasty (Noh et al., 2001), and after disrupting the auxin balance in flooded *R. palustris* (Cox et al., 2004). IAA accumulation does not occur in young leaves during waterlogging (Figure 8D), characterized by high *PIN4* and *PIN9* expression (Figure 8E – F), possibly sustaining auxin efflux (Figure 9).

Auxin transporters can be inhibited by flavonols (Kuhn et al., 2017). Flavonol overproducing mutants of Arabidopsis display a hyponastic phenotype, albeit in the leaf blade itself (Kuhn et al., 2011). In contrast, enhanced auxin transport in the *are* mutant in tomato reduces auxin accumulation (Maloney et al., 2014), which leads to a dampened epinastic response after waterlogging (Figure 6 & Figure 8). However, leaf epinasty in the *are* mutant can also be affected by changes in ROS metabolism and signaling.

Previously, it was proposed that auxins are redistributed towards the fast elongating side of the petiole (adaxial) during ethylene-induced epinasty in tomato (Lee et al., 2008) or during thermonasty (abaxial) in Arabidopsis (Park et al., 2019). In contrast, we observed a slight overall increase in DR5::GUS activity mostly at the abaxial side of older petioles during waterlogging.

Collectively, our data suggest that auxin accumulation by enhanced IAA production and reduced IAA export at the petiole base stimulates leaf epinasty in mature tomato leaves during waterlogging, while auxin transport is better sustained in young leaves, leading to stable IAA levels and thus a higher morphological plasticity during waterlogging (Figure 10).

### Ontogeny as an interface for regulating epinasty by ethylene and auxin crosstalk?

The concerted action of ethylene and auxins sets the scene for normal plant development (Van de Poel et al., 2015) and is often fine-tuned in a tissue-specific way (Muday et al., 2012), providing a framework for ontogenic regulation. Ethylene can either directly influence cell cycling and elongation (Dubois et al., 2018; Pierik et al., 2006) or change auxin homeostasis through modification of IAA production or transport. This dual action of ethylene could regulate the epinastic response in tomato, in spite of early reports on auxin-induced ethylene production regulating differential elongation in tomato petioles (Kazemi & Kefford, 1974; Stewart & Freebairn, 1969).

The auxin insensitive tomato mutant *diageotropica (dgt*) displays an epinastic response to both exogenous ethylene (Ursin & Bradford, 1989b) and waterlogging (Bradford & Yang, 1980), suggesting epinasty is primarily regulated by ethylene. On the other hand, reduction of ethylene signaling in the *Nr* mutant potentially affects cell elongation, but also enhances auxin transport in the hypocotyl, but not in roots (Negi et al., 2010). As a result, reduced epinasty in the *Nr* mutant might be the result of reduced ethylene sensitivity, but also of enhanced auxin transport (Figure 6; Negi et al., 2010) and disrupted auxin signaling (Lin et al., 2008). Altered ethylene signaling in this mutant causes development-dependent petiole bending (Figure 6), which might be due to ethylene and auxin responses. This is not surprising, given that ethylene and auxin signaling are integrated in regulatory hubs, such as *SlIAA3* (Chaabouni et al., 2009) and *SlERF.B3* (Liu et al., 2018) to establish tissue specific responses to both hormones. *SlIAA3,* a positive regulator of auxin responses in tomato petioles, is differentially expressed in petioles after ethylene treatment (Chaabouni et al., 2009), suggesting that it acts as an ethylene-mediated activator of auxin responses. The actual mechanism integrating ethylene and auxin responses in the petiole of tomato remains to be discovered.

## Acknowledgements

We thank the KU Leuven Greenhouse Core Facility for assistance in plant cultivation and Veerle Verdoodt for performing the ACC, MACC and ACO measurements. This work was funded by the Research Foundation Flanders with a FWO PhD fellowship (11C4319N; 1150822N; 1S02121N) to BG, JP and PM respectively, and a FWO research grant (G092419N) to BVDP; by a KU Leuven research grant (C14/18/056) to BVDP and a PhD-back-up grant (DB/17/007/BM) to BG. This work was also established in the framework of the RoxyCost action of the EU (CA18210).

## Author contribution

BG and BVdP designed the experiments, JP performed the RT-qPCR analyses and AMT activity assay, KVdB visualized and quantified cellular and macroscopic elongation of the petiole, VE assisted with physiological measurements, PM generated the EBS::GUS-GFP reporter line, ON supervised and designed the UHPLC-ESI-MS/MS protocol for auxin quantification, BG performed the experiments and analyzed the data, BG and BvDP wrote the manuscript. All authors read and approved the manuscript.

## Supporting Information

Supporting Information Method S1: Cloning and plant transformation

Supporting Information Method S2: Anthocyanin extraction

Supporting Information Method S3: Differential cell elongation of the abaxial – adaxial petiole side

Supporting Information Table S1: Primers used for the qRT-PCR in this study

Supporting Information Table S2: Primers used for the tomato transformation in this study

Supporting Information Fig. S1: Petiole morphology and anatomy of the different leaves of tomato plants in the eighth leaf stage.

Supporting Information Fig. S2: Daily transpiration of tomato during and after a waterlogging treatment of 96 h.

Supporting Information Fig. S3: Effect of waterlogging (48 h) on biomass production in leaves of different ages.

Supporting Information Fig. S4: Effect of waterlogging (96 h) on biomass production in leaves of different ages.

Supporting Information Fig. S5: Petiole ethylene production after sampling.

Supporting Information Fig. S6: GUS expression in petioles of different ages in an *EBS::GUS* reporter line.

Supporting Information Fig. S7: Auxin responses and biosynthesis in leaves and petioles of different ages.

Supporting Information Fig. S8: Effect of inhibition of auxin transport (TIBA) on leaf epinasty during waterlogging.

Supporting Information Fig. S9: Morphological effects of a local TIBA application with lanolin paste on the petiole.

Supporting Information Fig. S10: DR5::GUS staining pattern in petioles treated with a 0.5 – 1 cm ring of TIBA in lanolin paste at the petiole base.

Supporting Information Fig. S11: Concentration of anthocyanins, natural inhibitors of auxin flows, in the different leaves.

## Notes

### Competing Interest Statement

The authors have declared no competing interest.

