## Supporting Information for "Leaf ontogeny steers ethylene and auxin crosstalk to regulate leaf epinasty during waterlogging of tomato"

Supporting Information Method S1: Cloning and plant transformation

Supporting Information Method S2: Anthocyanin extraction

Supporting Information Method S3: Differential cell elongation of the abaxial – adaxial petiole side

Supporting Information Table S1: Primers used for the qRT-PCR in this study

Supporting Information Table S2: Primers used for the tomato transformation in this study

Supporting Information Fig. S1: Petiole morphology and anatomy of the different leaves of tomato plants in the eighth leaf stage.

Supporting Information Fig. S2: Daily transpiration of tomato during and after a waterlogging treatment of 96 h.

Supporting Information Fig. S3: Effect of waterlogging (48 h) on biomass production in leaves of different ages.

Supporting Information Fig. S4: Effect of waterlogging (96 h) on biomass production in leaves of different ages.

Supporting Information Fig. S5: Petiole ethylene production after sampling.

Supporting Information Fig. S6: GUS expression in petioles of different ages in an *EBS::GUS* reporter line.

Supporting Information Fig. S7: Auxin responses and biosynthesis in leaves and petioles of different ages.

Supporting Information Fig. S8: Effect of inhibition of auxin transport (TIBA) on leaf epinasty during waterlogging.

Supporting Information Fig. S9: Morphological effects of a local TIBA application with lanolin paste on the petiole.

Supporting Information Fig. S10: DR5::GUS staining pattern in petioles treated with a 0.5 – 1 cm ring of TIBA in lanolin paste at the petiole base.

Supporting Information Fig. S11: Concentration of anthocyanins, natural inhibitors of auxin flows, in the different leaves.

### Supporting Information Method S1: Cloning and plant transformation

The *EBS::GFP-GUS* reporter line was generated in the Ailsa Craig background. The *EBS::GUS-GFP* fusion plasmid was constructed with an *EBS* sequence cloned from an Arabidopsis *EBS::GUS* reporter line (NACS CS69047), described in Stepanova et al. (2007), and fused to the *GFP-GUS* sequence, cloned from the *pKGWFS7.1* vector, in the *pCAMBIA1302* destination vector. All PCR products were cloned using Gibson assembly after digestion with *BstE*II and *Sal*I. The *pCAMBIA EBS::GFP-GUS* destination vector was transformed into tomato using agrobacterium-mediated cotyledon transformation following a modified protocol by Van Eck et al. (2019). Primers are listed in Supplemental Table 2.

### Supporting Information Method S2: Anthocyanin extraction

Anthocyanin levels were measured spectrophotometrically, following the protocol described by Maloney et al. (2014b). After extraction, the absorbance was measured at 530 nm and corrected with the absorbance at 675 nm to quantify anthocyanin content.

### Supporting Information Method S3: Differential cell elongation of the abaxial – adaxial petiole side

We quantified the relative cell elongation by comparing cell length distribution quantiles (sample population of cells sorted by their length and divided in 10 % groups based on their cell length) in 0.5 cm zones on the adaxial and abaxial side of the petiole (Figure 1E and Supporting Information Method S3 Figure). Cells at the higher end of the distribution, thus already large cells, became proportionally larger in petioles of waterlogged plants. Differences between these cell quantiles ranged from 0 to 100 µm and more after the treatment, depending on the position along the petiole as well as leaf age. In addition, leaf development seemed to limit the extent of the differentiation between adaxial and abaxial epidermis cells. In leaves 5 and 7, for example, elongation of the adaxial epidermis cells is more prominent in a larger part of the petiole, while in leaf 3 it seems mostly confined to the region closest to the stem. The shift function also revealed that in control conditions, cells at the abaxial side elongated relatively more than those at the adaxial side, reflected by a higher upper size limit for these cells, mostly in older leaves (1 – 4). As a result, the differential elongation across the petiole was respectively larger during waterlogging conditions, which in turn led to downward bending of the petiole. This mechanism is depicted in Figure 1G for the zone closest to the petiole base (zone 1) in leaf 3. In this example, 10 %, 40 %, 70 % and 90 % of the total cell population are smaller than the cells depicted in the model.

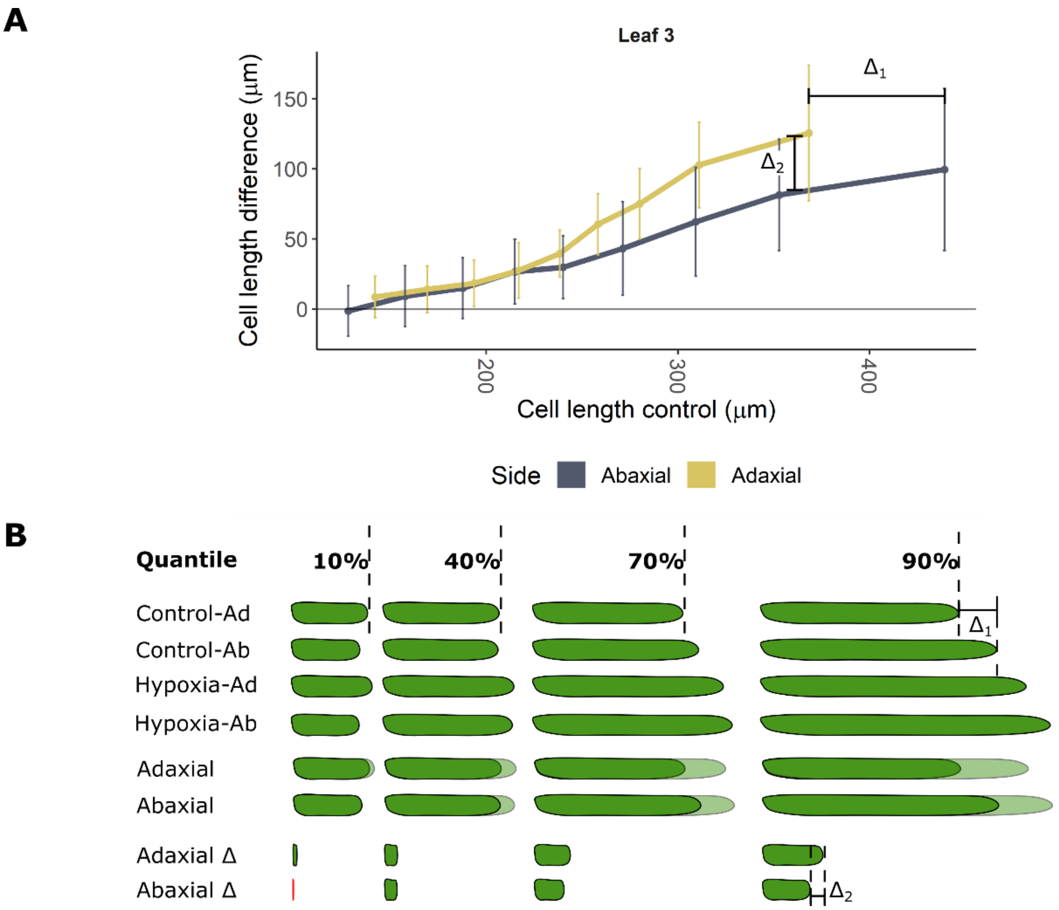

Supporting Information Method S3 Figure: Schematic representation of the shift function cell elongation in the first 0.5 cm (zone 1) of the adaxial and abaxial sides of the petiole of leaf 3 during waterlogging. (A) Example of the quantile differences between cells in waterlogging and control conditions for leaf 3 after 48 h of waterlogging. (B) Model of individual cells at the adaxial and abaxial side of the petiole. The model depicts the average cell length for the quantile thresholds (10 %, 40 %, 70 % and 90 % of all cells are smaller than the cell size depicted here) at each side of the petiole, the superposition of these cells and the differences between abaxial and adaxial cell lengths of control and treated plants. $\Delta_1$ and $\Delta_2$ are the differences between the abaxial and adaxial petiole cell length and between adaxial and abaxial elongation of treated and control plants respectively. Lines in (B) represent average leaf angles +/- the confidence interval (90 %).

Supporting Information Table S1: Primers used for the qRT-PCR in this study

| *Gene* | Solyc | Forward | Reverse |
|---|---|---|---|
| *ACO1* | Solyc07g049530 | TTG CGC CAT CTT CCT ACT TC | CTC CTC AGC CAA TTT CTC CA |
| *ACO2* | Solyc12g005940 | TGC ACA TAC TGA TGC TGG TG | TCG ACC GTC TTT GAG GAG TT |
| *ACO4* | Solyc02g081190 | CTG GGG TTT CTT TGA GCT TG | TCC TTT ACT TGC CAC CAT TTC T |
| *ACO5* | Solyc07g026650 | ACC GAG TGA TGG CTG AAA AG | TAA TCC TGA AAG CGG AGG TG |
| *ACO6* | Solyc02g036350 | AAC TGG GGA TTC TTT GAG GTG | TTA CTT GCC ACC ATT TCC TTG |
| *ACO7* | Solyc06g060070 | TGA TGC TGG TGG TGT CAT TT | CAC GAT GGC GTT AGG GAT AG |
| *MoOx12* | Solyc08g068160 | ACC CCT GTA CTT GAC GTT GG | ATT CCA CCG TGT GAT GCT TT |
| *PIN4* | Solyc05g008060 | GTT CAG AGT TCC CCG AGG TT | ATT TGG CTG CTG AGG TTT TG |
| *PIN9* | Solyc10g078370 | TGT CTT TAG GGG AGG AGG TG | CGG TCA TTT CCA TTC AGG TT |
| *ACT* | Solyc03g078400 | CAA CAG AGA GAA GAT GAC CCA | ACC AGA GTC CAA CAC AAT ACC A |
| *EF1a* | Solyc06g005060 | ACA GGC GTT CAG GTA AGG AA | TGG AGG GTA TTC AGC AAA GG |
| *PP2A* | Solyc05g006590 | TGG TTG CTT TGA AGG TTC GT | TTG GCA TTT CCA TAC TTC CTC |
| *RPL2* | Solyc10g006580 | GAG GGC GTA CTG AGA AAC CA | TCA TAG CAA CAC CAC GAA CC |

Supporting Information Table S2: Primers used for the tomato transformation in this study

| *Insert* | *Forward* | *Reverse* |
|---|---|---|
| at_EBS | ggggacaagtttgtacaaaaaagcaggctTGCGAAGGATAGTGGGATTG | ggaccactttgtacaagaaagctgggtGATCTAGTAACATAGATGACACCGC |
| GFP-GUS | tggagaggacagggtatcaaATGGTGAGCAAGGGCGAG | cgatcggggaaattcgagctgTCATTGTTTGCCTCCCTGC |
| EBS | tacccggggatcctctagagACAGCTATGACCATGATTAC | ctcctcgcccttgctcaccatTTGATACCCTGTCCTCTC |

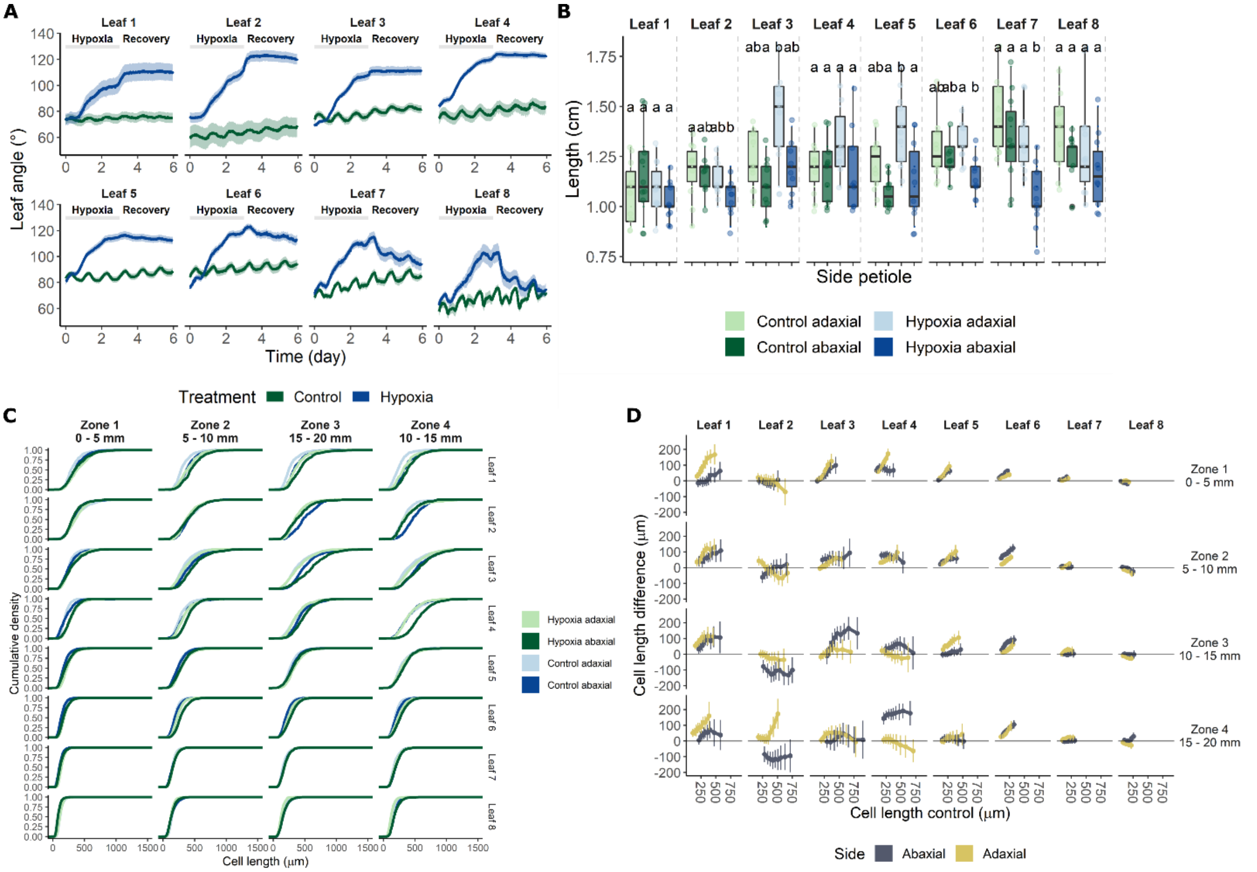

Supporting Information Fig. S1: Petiole morphology and anatomy of the different leaves of tomato plants in the eighth leaf stage. (A) Leaf angle dynamics during and after a 72 h waterlogging treatment (n = 10). (B) Length of the abaxial and adaxial sides (n = 10) of the first cm of the petiole after 48 h of treatment. (C) Empirical cumulative cell length distributions and (D) differences in epidermal cell length distributions (n = 2 – 631 cells per treatment and zone for the adaxial and abaxial side per plant; 6 – 7 plants per treatment) between 48 h waterlogging and control conditions. The different zones in (D) refer to regions of the petiole of approximately 5 mm, starting at the petiole base (zone 1) and stretching out 20 mm along the petiole (zone 4). Significant differences are indicated with letters (α = 0.05).

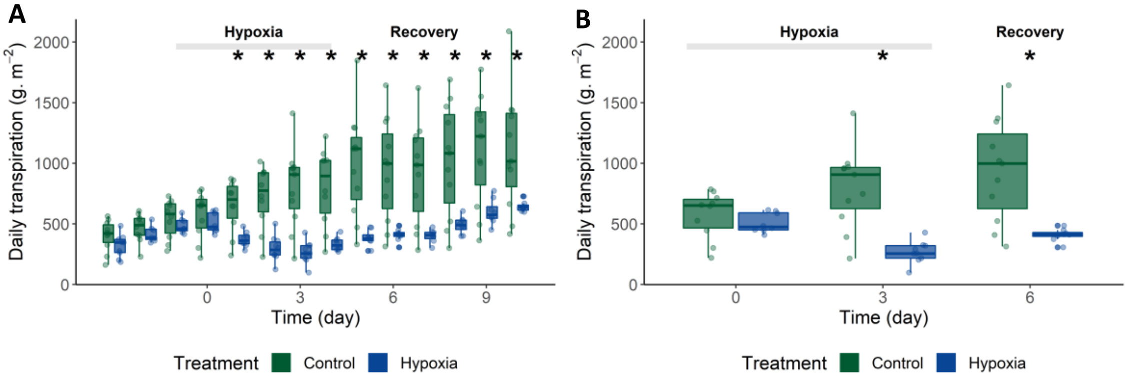

Supporting Information Fig. S2: Daily transpiration of tomato during and after a waterlogging treatment of 96 h. Transpiration for (A) the entire period and for (B) the start and end of the treatment and the 3rd day of recovery. Significant differences are indicated with an asterisk (α = 0.05).

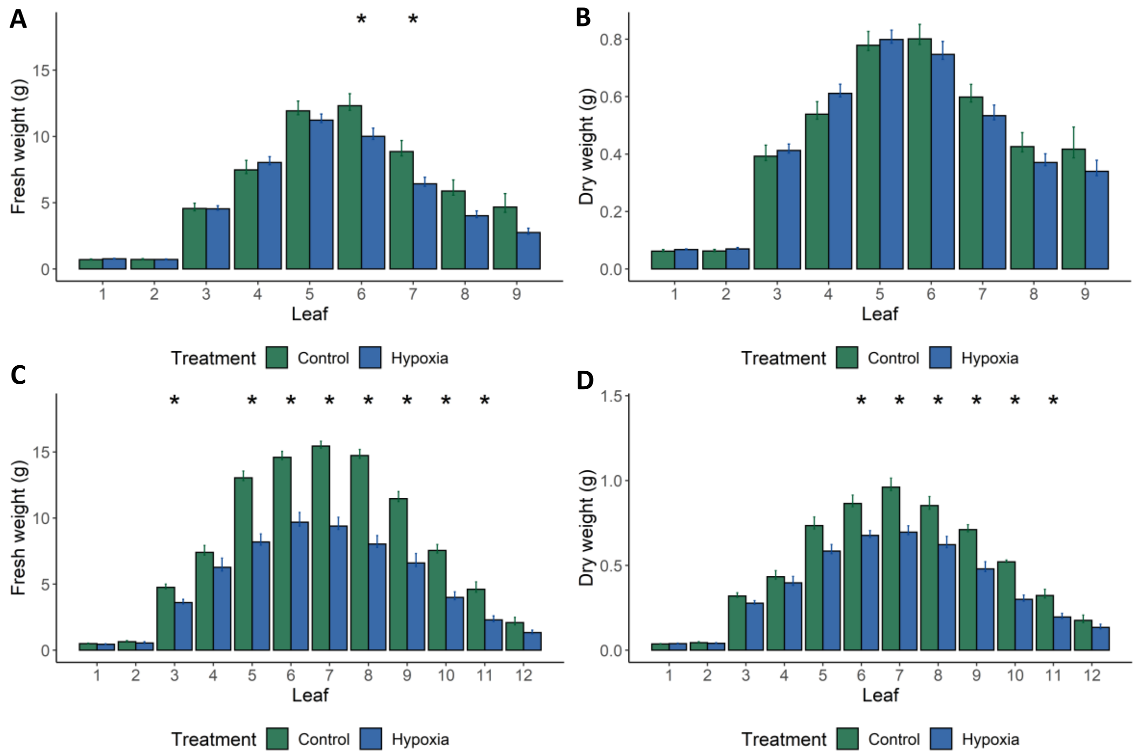

Supporting Information Fig. S3: Effect of 48 h of waterlogging on biomass production of leaves of different ages. (A) Fresh weight and (B) dry weight after a 48 h waterlogging treatment and (C – D) an additional reoxygenation of one week. Significant differences are indicated with an asterisk (α = 0.05).

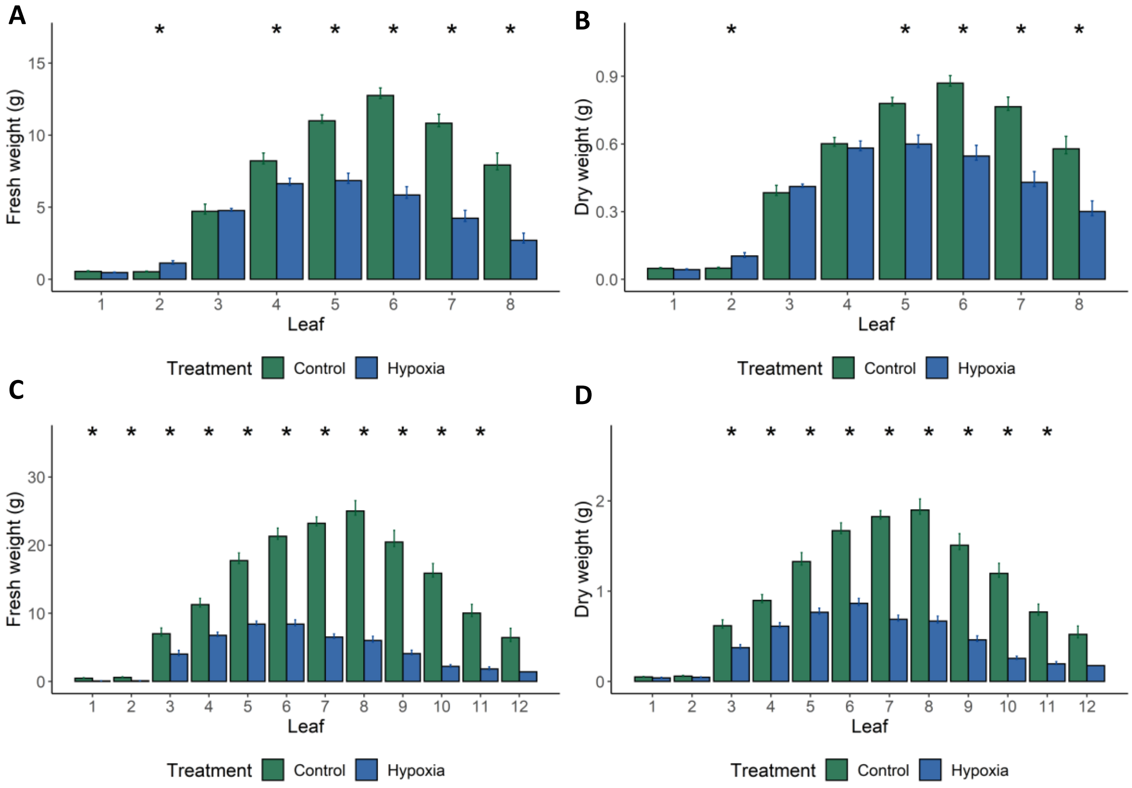

Supporting Information Fig. S4: Effect of 96 h of waterlogging on biomass production of leaves of different ages. (A) Fresh weight and (B) dry weight after a 96 h waterlogging treatment and (C – D) an additional reoxygenation of one week. Significant differences are indicated with an asterisk (α = 0.05).

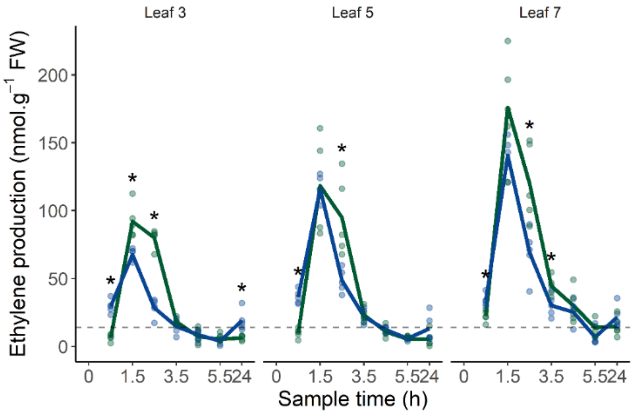

Supporting Information Fig. S5: Petiole ethylene production profiles during 24 h after sampling. Significant differences are indicated with an asterisk (α = 0.05).

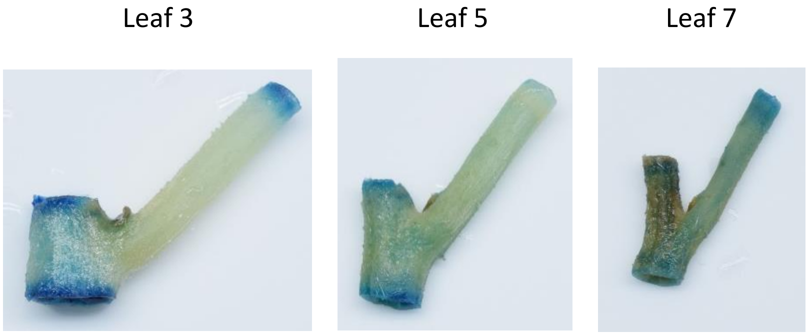

Supporting Information Fig. S6: GUS expression in petioles of different ages of an *EBS::GUS* reporter line after waterlogging for 48 h.

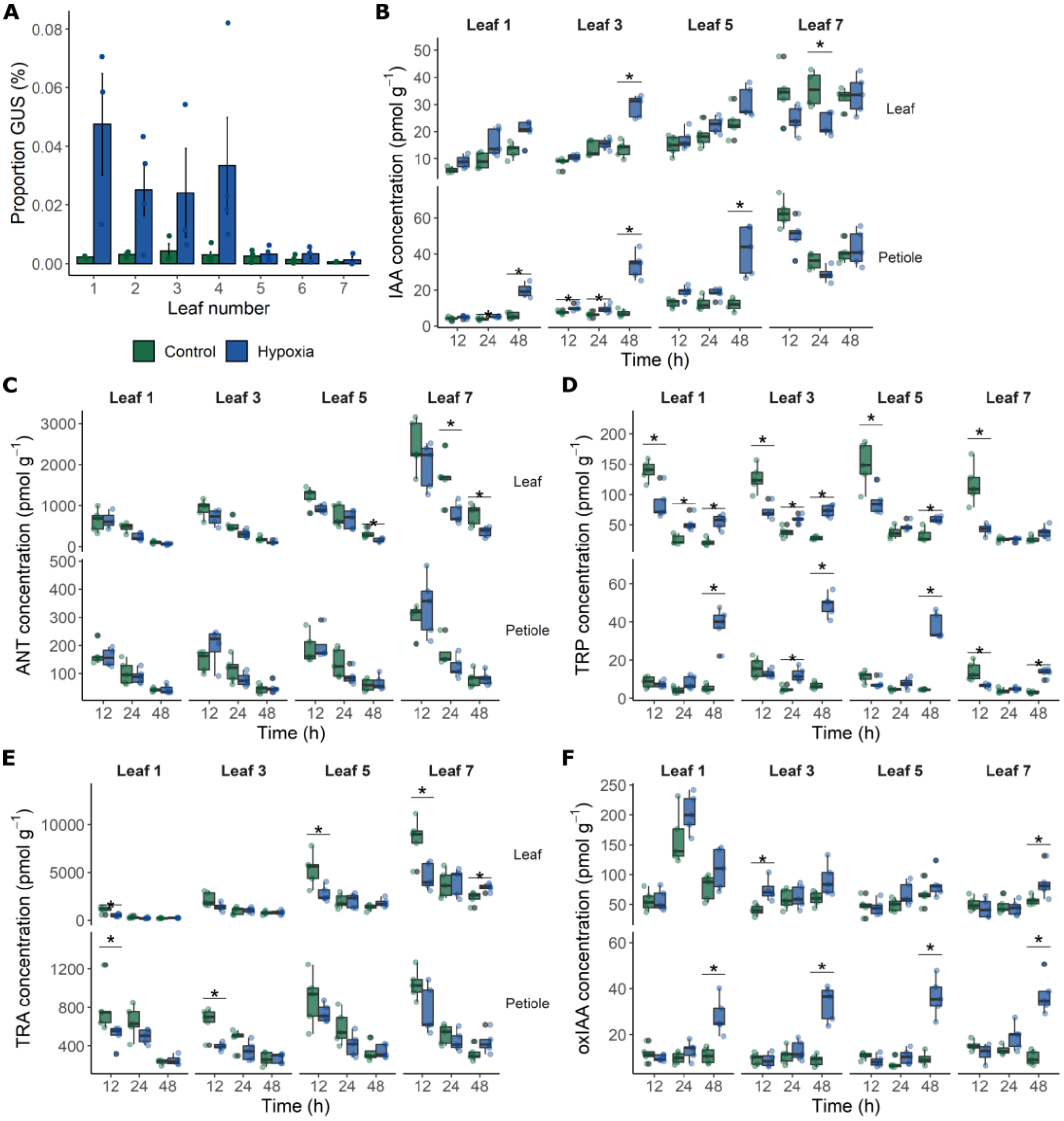

Supporting Information Fig. S7: Auxin responses and biosynthesis in leaves and petioles of different ages. (A) Quantification of GUS expression in petiole cross sections after 48 h of waterlogging. (B – F) Auxin metabolism in tomato leaf and petiole tissue during a waterlogging treatment (n = 5). Concentration of (B) IAA, (C) ANT, (D) TRP, (E) TRA, and (F) oxIAA. Significant differences are indicated with an asterisk (α = 0.05).

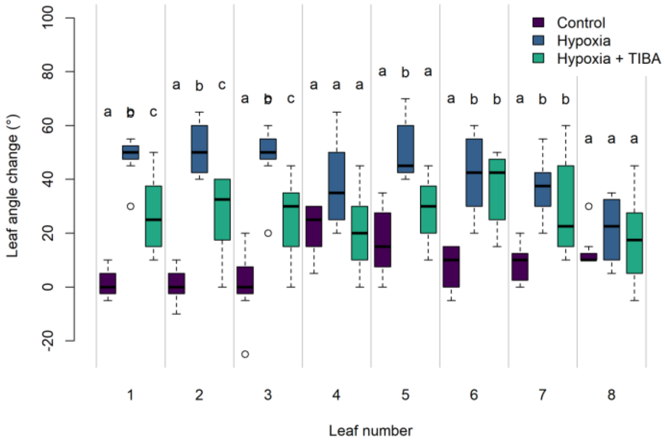

Supporting Information Fig. S8: Effect of inhibition of auxin transport (TIBA) on leaf epinasty during waterlogging. Leaf angle change after 48 h of waterlogging with and without a foliar application of TIBA the night before the start of the treatment. Significant differences are indicated with letters (α = 0.05).

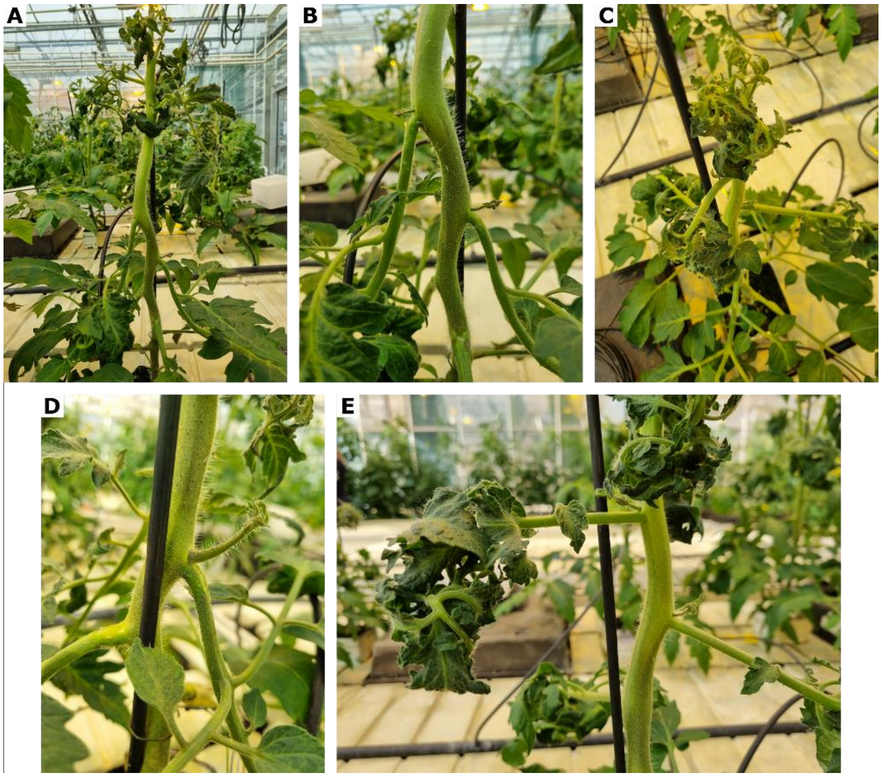

Supporting Information Fig. S9: Morphological effects of a local TIBA application with lanolin paste on the petiole. (A – B) Effect on leaf morphology of (A) a control plant, characterized by (B) strong epinasty of treated leaves and (C) strong curling of untreated, young leaves. (D – E) Effect on morphology of a plant after 72 h of waterlogging in combination with the TIBA application. (D) Epinastic bending of petioles treated with TIBA and (E) straight petiole/rachis morphology and curled leaflets of untreated leaves.

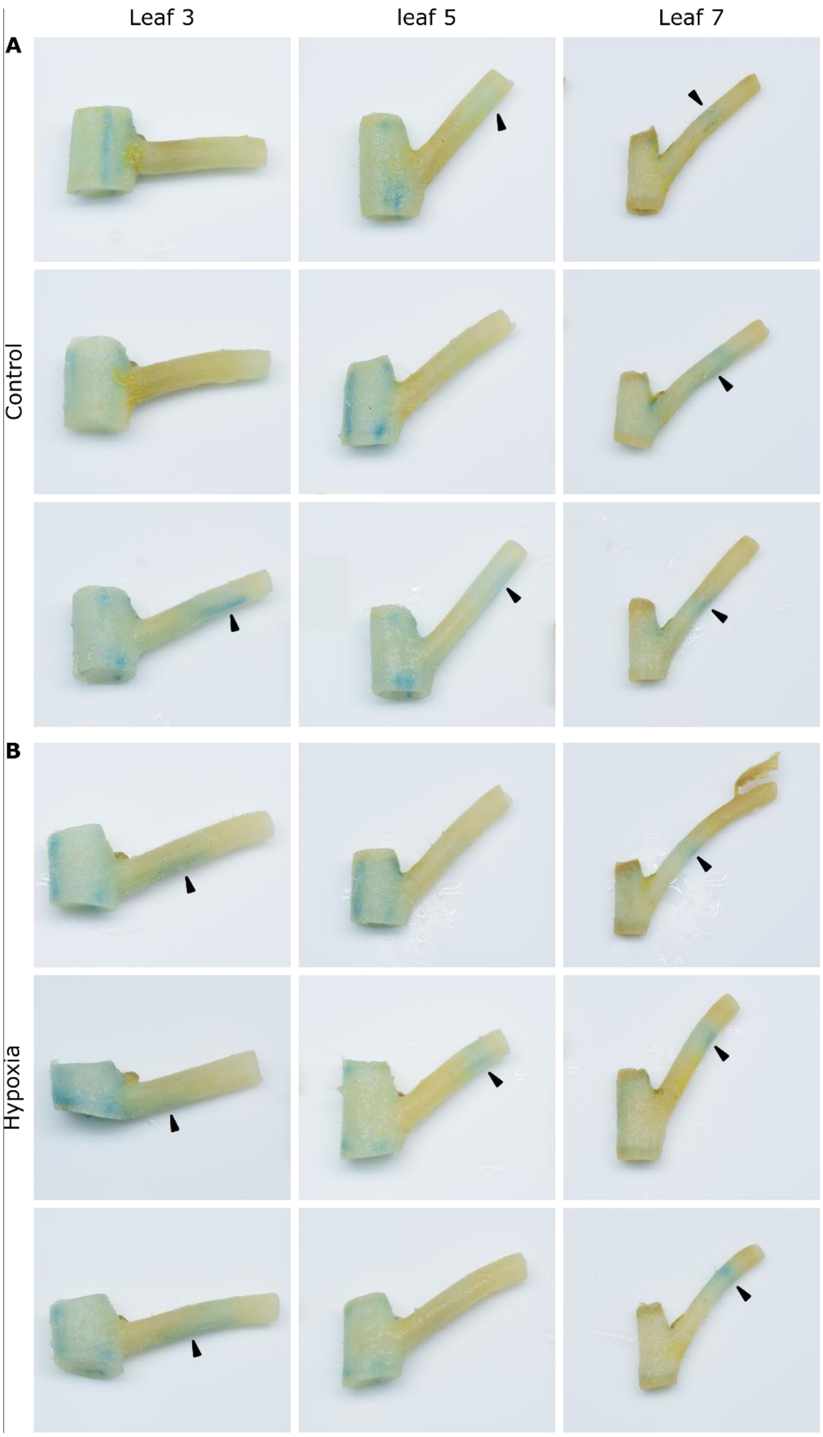

Supporting Information Fig. S10: DR5::GUS staining pattern in petioles treated with a 0.5 – 1 cm ring of TIBA in lanolin paste at the petiole base. Petioles of leaf 3, 5 and 7 of (A) 3 control plants and of (B) 3 plants after a 24 h waterlogging treatment.

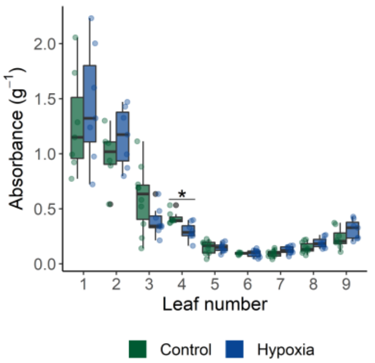

Supporting Information Fig. S11: Concentration of anthocyanins, natural inhibitors of auxin flows, in the different leaves. Anthocyanin levels (n = 7 – 10) were derived from spectrophotometric analyses. Significant differences are indicated with an asterisk (α = 0.05).
